# Compensatory tRNA Modification by DUS3L Confers Resistance to METTL1 Loss in Oesophageal Cancer

**DOI:** 10.1101/2025.10.13.682082

**Authors:** Helena Santos-Rosa, Jonathan L. Price, Georgia Tsagkogeorga, Alexandra Sapetschnig, Bianca Pierattini, David Jordan, Paula Milan-Rois, Almudena Serrano-Benitez, Katherine Stott, Namshik Han, Tony Kouzarides, Eric A. Miska

## Abstract

tRNA N⁷-methylguanosine (m⁷G46), installed by the METTL1/WDR4 methyltransferase, stabilizes the tRNA pool supporting high proliferative rates. Accordingly, many cancer cell lines are vulnerable to METTL1 loss, underscoring its potential as a therapeutic target. We identify a subset of cell lines insensitive to METTL1 depletion and use oesophageal lines as a model to explore the mechanism. The METTL1-resistant adenocarcinoma line OE33 preserves tRNA balance upon m⁷G46 deficiency by elevating dihydrouridylation at position 47 (D47). Depletion of DUS-L enzymes that catalyse D16/D17/D20/D47 on tRNAs sensitises OE33 to METTL1 loss. Concomitant loss of dihydrouridylation and m⁷G46 destabilizes the canonical L-shaped tRNA structure, disrupts coaxial helix stacking and anticodon-loop presentation, and reduces cellular fitness. Notably, METTL1-sensitive cell lines lack this compensatory pathway due to the deleterious effects of increasing their intrinsically low DUS3L levels. These findings reveal a DUS-L–mediated buffering mechanism that supports tRNA homeostasis in METTL1-resistant cancers and highlight DUS3L abundance as a potential biomarker for predicting METTL1 sensitivity. Moreover, they suggest that combined targeting of DUS-L and METTL1 may provide a therapeutic strategy for overcoming resistance in cancers, exemplified in cell lines such as OE33.

**Highlights:**

- The oesophagus cancer line OE33 shows no proliferative defect upon METTL1/WDR4 loss, despite complete lack of m^7^G46 on its tRNA.
- OE33 coordinates codon usage and abundance of cognate tRNAs in the absence of METTL1/WDR4; the METTL1/WDR4 sensitive line, OE21, fails to do so.
- Depleting dihydrouridine synthases (DUS-L) renders OE33 sensitive to METTL1/WDR4 loss.
- OE33 responds to lack of METTL1/WDR4 activity by upregulating DUS3L and its activity (D47) on tRNA; the sensitive line OE21 fails to do so.
- METTL1-sensitive lines have intrinsically low DUS3L and cannot sustain higher levels.
- DUS3L reveals as a potential biomarker for METTL1 sensitivity.

## Introduction

RNAs are decorated with >150 post-transcriptional modifications occurring co-and post-transcriptionally at structural and functional sites such as caps, splice-junctions, 5′-and 3′-UTRs or coding regions. These modifications fine-tune most aspects of cell biology affecting RNA structure, half-life, subcellular localization, translatability and interactions with other RNAs, DNAs or proteins ^1,2^.

Unsurprisingly, an imbalanced dosage of RNA-modifying enzymes is pathogenic. The overexpression of RNA writers, such as METTL3, NSUN2, PCIF1/CAPAM, and erasers, like FTO, reprogram oncogenic, metabolic and immune pathways, whereas loss-of-function mutations in tRNA, rRNA or editing enzymes (*NSUN2, TRMT10A, DKC1, ADAR1*) trigger neurodevelopmental delay, diabetes, marrow failure and interferonopathy^3^.

Transfer RNA (tRNA) is the most densely modified RNA, with the highest incidence of events in the anticodon loop and at the D/T junction, reflecting their key roles in decoding accuracy and stable 3-D folding respectively^4,5^. One of these modifications is the evolutionarily conserved N7-methylation of guanosine at position 46 in the variable loop of the tRNAs. Upon methylation, G46 gets positively charged at physiological pH, stabilizing tertiary interactions between T/D loops that are critical to provide the canonical tRNA’s L-folded shape^6–8^. Work in *S. cerevisiae* showed that the lack of several post-transcriptional modifications leads to destabilization and degradation of mature tRNAs via the Rapid tRNA Degradation pathway (RTD), where Rat1/Xrn1 5′→3′ exonucleases dismantle faulty mature molecules ^9–12^. Recent work in mammal advocates for a conserved mechanism^13,14^. The enzymatic activity responsible for m^7^G46 was initially identified in budding yeast as consisting of the methyltransferase Trm8 and its cofactor Trm82^7^. Recessive mutations in the mammalian orthologues, *METTL1* and *WDR4*, result in microcephalic primordial dwarfism and Galloway-Mowat syndrome^15–16^. Conversely, copy-number gains or super-enhancer activation of *METTL1* is widespread in leukaemia and solid tumours, correlating with vascular invasion, increased proliferation and chemoresistance. These facts make the METTL1/WDR4 complex a compelling therapeutic target in oncology ^15–19^ and initial efforts towards METTL1 inhibition by small molecules have been disclosed with promising results^20,21^. Yet, some malignancies seem to exhibit intrinsic resistance to the lack of METTL1/WDR4^22^ and acquired resistance mechanisms need to be foreseen.

Here, we investigate the molecular pathways underlying resistance to METTL1 loss, using oesophagus cancer cell lines as a model system. By leveraging a CRISPR-Cas9 synthetic lethal screen in the resistant OE33 cell line, we identify several members of the DUS-L family of dihydrouridine synthases as a molecular network that support proliferation in the absence of METTL1/WDR4 activity. Furthermore, our data uncovers the specific crosstalk between METTL1 and DUS3L activities (m^7^G46 and D47 respectively) pointing towards a structural compensatory mechanism that naturally resistant, but not sensitive, cancer cell lines use to adjust the cellular tRNA pool to their needs. Hence, our work reveals the DUS-L family as an escape route for rational combination therapies, that would expand the panel of treatable malignancies and reinforce METTL1 inhibitors into durable anticancer agents.

## RESULTS

### Heterogeneous response of cancer cells to METTL1 down-regulation

Over the past decade, numerous studies have shown that depletion of the tRNA m^7^G46 methyltransferase METTL1 attenuates the proliferative potential of a wide spectrum of cancer cell lines^18,19^. Large scale CRISPR screens catalogued in DepMap broadly corroborate this dependency, while also revealing several outliers^22^. To investigate the molecular basis of such METTL1 independence, we performed siRNA-mediated knockdown of METTL1 in a random panel of lines representing diverse tumour types. Immunoblotting confirmed robust loss of METTL1 protein in all cell lines (Figure S1A). As anticipated, our results show that METTL1 silencing impairs proliferation in several lines (Fig. 1A; Figure S1A and 1B). Yet, a subset exhibited no growth defect or even a mild proliferative advantage upon METTL1 depletion (Figure 1A; Figure S1A and S1B). This heterogeneous response was surprising considering that METTL1/WDR4 activity (N7-methylguanosine) is known to play a very important role in the proper folding and stability of the cellular tRNA pool. To characterise these responses further, we chose oesophagus cancer as our model system given the availability of a sensitive (OE21, oesophagus squamous cell carcinoma) and a resistant (OE33, oesophagus adenocarcinoma) cell line (Figure 1A, and Figure S1A). Competition assay by CRISPR-Cas9 technology to knockout *METTL1*, phenocopied the results obtained by METTL1 siRNA (Figure S1C), corroborating the METTL1 sensitivity of OE21 and resistance of OE33. Of note, down regulation of METTL1 resulted in the expected instability of the cofactor WDR4 in both cell lines, pointing to the fact that the interdependency of enzyme/cofactor is common to both, despite their different proliferative responses to the lack of METTL1 (Figure S1D).

**Figure 1.**
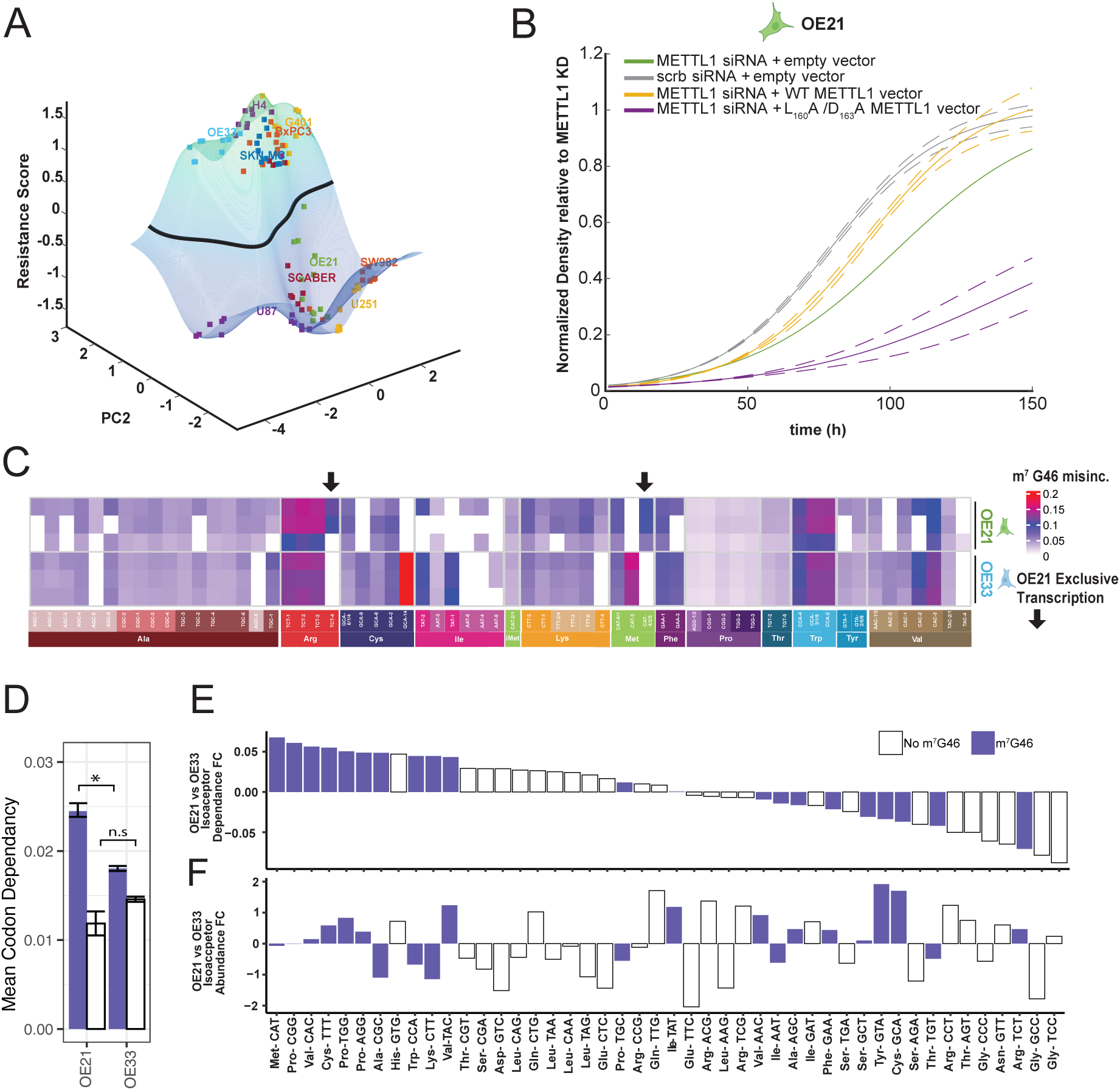
Characterization of METTL1 sensitive (OE21) and resistant (OE33) Oesophagus cancer cell lines. A. Proliferative Dependency on METTL1 across cancer cell lines. Each point represents the first and second principal component in the X and Y axis. The PCA was performed on the parameters from the logistic fits of each growth curve. A kernel SVM classifier was applied to the PCA data to find the best separation between resistant and sensitive cell lines. The SVM classifier score represents a “resistance” score, where high values in Z correspond to resistance and low values to sensitivity. H4: Astrocytoma, G401: Kidney Rhabdoid, BXPC3: Pancreatic Adenocarcinoma, OE33: Oesophagus Adenocarcinoma, SKN-MC: Neuroepithelioma, OE21: Oesophagus Squamous Carcinoma, SCABER: Bladder Squamous Carcinoma, SW982: Synovial Sarcoma, U251: Astrocytoma, U87: Glioblastoma. B. Wild-type METTL1, but not catalytic mutant, rescues OE21 proliferation defects. Stable multiclonal OE21 populations expressing either wildtype METTL1, METTL1 L_160_AD_163_A catalytic mutant (both alleles siRNA resistant) or empty vector, were transfected with 2.5nM METTL1 siRNA and non-targeting scramble control.18h post-transfection the medium was changed, and proliferation was followed for 150h by Incucyte. The results represent 3 independent experiments (n=3), each consisting of biological triplicates. Dotted lines represent 95% CI. C. OE21 and OE33 specific distribution of m⁷G46 across tRNA isodecoders. Heatmap of m⁷G46 signal in OE21 and OE33 cells obtained by sodium borohydride–dependent misincorporation mapping (mim-tRNA-seq). Columns correspond to individual tRNA isoacceptors (grouped by amino-acid identity; anticodons shown below). Colour indicates the misincorporation ratio at position 46 (non-reference calls/total reads at G46), which serves as a proxy for m⁷G46 occupancy (scale at right; higher values = higher m⁷G46). Only isoacceptors with sufficient read coverage are displayed; white tiles indicate no detectable signal. Black arrows denote isodecoders expressed in only one cell line, as indicated by POLR3A occupancy at the corresponding loci. D. Mean Codon dependency for methylated and unmethylated isoacceptors, *t* (2) = 8.9, p = 0.01. E. Codon-level isoacceptor usage in OE21 *versus* OE33. Fold change (FC) in codon dependence derived from RNA-seq–based codon usage analysis comparing OE21 and OE33 cells. Bars represent codons decoded by tRNAs carrying m⁷G46 modifications (purple) or unmodified tRNAs (white). Aminoacid identity is shown under each anticodon. (n=3) F. tRNA isoacceptor abundance in OE21 *versus* OE33. Fold change (FC) in codon dependence derived from tRNA-mim-seq analysis comparing OE21 and OE33 cells. Bars represent tRNAs isoacceptors carrying m⁷G46 modifications (purple) or unmodified tRNAs (white). Aminoacid identity is shown under each anticodon. (n=3)

To prove that the growth defect observed in OE21 was owed to METTL1 enzymatic activity, we performed the rescue experiment by co-expression of WDR4 and siRNA resistant *METTL1* cDNAs, either wild-type or catalytic dead (*METTL1 L_160_AD_163_A*)^13^ (Figure S1E). Wild-type and mutant alleles were expressed at the same level as assessed by western blot with specific antibody (Figure S1F, *tMettl1). We could confirm that wild-type METTL1 (Figure 1B, yellow) but not the catalytic mutant (Figure 1B, purple) rescued the growth of OE21 upon METTL1 siRNA (Figure 1B, green), revealing the lack of enzymatic activity as responsible for the phenotype. These data establish oesophageal carcinoma as a tractable model to elucidate context specific mechanisms that bypass METTL1 mediated tRNA m^7^G46 modification.

### tRNA loci-usage and codon-usage is consistent between METTL1 sensitive and resistance cell lines

To investigate the reasons for the METTL1 response of the sensitive (OE21) and the resistant (OE33) cell-lines, we first mapped their specific m^7^G46 methylome by chemical treatment with Na-borohydride followed by RNA-seq (mim-tRNAseq). As shown in Figure 1C, the repertoire of METTL1/WDR4 substrates largely overlapped the 17–22 isodecoders previously reported to harbour m^7^G46 in other studies^23^. Although the m^7^G46 targets largely coincide between the sensitive and resistant cell-lines, a distinct subset of tRNAs was uniquely modified in each one.

Notably, different members of the tRNA-Met-CAT isodecoder family were preferentially modified in each cell line (tRNA-Met-CAT2/4/5 in OE21 versus tRNA-Met-CAT3 in OE33). Moreover, tRNA-Cys-GCA14 was heavily methylated in OE33, whereas the tRNA-Arg-CTC4, described as “oncogenic-driver”^14^, lacked detectable m^7^G46 in this line (Figure 1C).

To investigate whether the absence of specific tRNA species in each cell line’s m⁷G46 methylome was due to a lack of METTL1 activity on those substrates or simply reflected their transcript abundance, we performed POLR3A chromatin immunoprecipitation (RNA POL III-ChIP). This is an established *proxy* for tRNA expression because POL III does not pause at gene promoters and thus unambiguously maps the genomic location and transcription state of each tRNA loci^24^. We validated the specificity of our ChIPs by Q-PCR using primers to Leu-tRNA and 5S-ribosomal RNA, both POLIII transcripts (Figure S1G). We did not find significant difference in loci usage between OE21 (sensitive) and OE33 (resistant) except for a dozen (out of more than 500 tRNA loci across the genome). Among the few exceptions are the “oncogenic” tRNA-Arg-CTC4 and tRNA-Met-CAT2/4/5 loci which are expressed OE21 but not OE33, explaining their absence in the methylome of the later one (Figure 1C and Figure S1H, black arrows). Overall, members of different isodecoder families harbouring the consensus for METTL1/WDR4 activity (‘“RAGGU”) but absent in our m^7^G46 mim-tRNAseq datasets where simply very low or not expressed. Yet, at isoacceptor and isotype levels tRNA transcripts show no difference between the OE21 and OE33 (Figure S1H). We conclude that the specific m⁷G46 methylome is determined by the tRNA transcriptional state together with few cell-line METTL1/WDR4 substrate preferences. Overall, at the isoacceptor and isotype levels, the methylomes of OE21 and OE33 show no significant differences—except for Cys— as it was the case for the tRNA expression profile (Figure S1I).

We then examined codon usage to assess potential bias between the two cell lines. Specifically, we asked whether OE21 transcriptome relies more heavily than OE33 on codons decoded by m⁷G46-modified tRNAs. The analysis revealed a significant difference between OE21 and OE33 pointing to a greater dependency on modified tRNAs (Figure 1D and 1E). Over the past decade, studies have documented a significant association among codon usage, tRNA abundance, and the expression of cell-proliferation genes in mammals^25–29^. We reasoned that if the preferential use of codons decoded by m⁷G46-modified tRNAs was a key determinant of the METTL1 sensitivity of OE21 cells, it may be reflected in their abundance. However, OE21 showed no overall enrichment of m⁷G46-modified tRNAs —compared with OE33— to which attribute their differential response to METTL1 loss (Figure 1F).

Taken together, we found no global differences in loci usage, m⁷G46 tRNA profiles, or tRNA abundance that could account for the distinct METTL1 dependence of OE21 and OE33 cells.

### OE33 maintains tRNA homeostasis despite m⁷G46 deficiency

To further investigate potential differences, we set out to produce single cell clones deficient in METTL1/WDR4 activity from resistant (OE33) and sensitive (OE21) cell lines. We generated OE33 *METTL1* knockout clones by transfecting guide RNAs against *METTL1* exon1 and exon2, without any selection step (Figure S2A and Methods). Genotyping showed an almost perfect Mendelian distribution (19 % WT/WT, 28.6 % KO/KO, 52.4 % WT/KO), corroborating that loss of METTL1 does not confer a proliferative disadvantage. Immunoblotting confirmed the absence of METTL1 protein in the knockout clones and, in addition, revealed a marked reduction in WDR4 levels (Figure 2A). Proliferation assays demonstrated that *METTL1*KO clones phenocopied METTL1 siRNA knockdown in OE33, displaying growth rates that were unchanged or slightly higher than those of the wildtype single cell control produced with non-targeting guide RNAs and the parental cell line (Figure S2B). Despite their normal fitness, all *METTL1*KO clones completely lacked m^7^G on their tRNA, in stark contrast to the wildtype clone and parental cells, as assessed by North-western blotting with a m^7^G antibody (Figure 2B). Modification-induced-misincorporation (mim-tRNAseq) profiling proved that deletion of *METTL1* in OE33 results in complete abrogation of m^7^G46 from tRNAs, confirming the above (Figure 2C). Together, these data indicate that no alternative enzyme compensates for METTL1 loss in OE33 and that tRNA m^7^G46 is dispensable for proliferation in this cell line.

**Figure 2.**
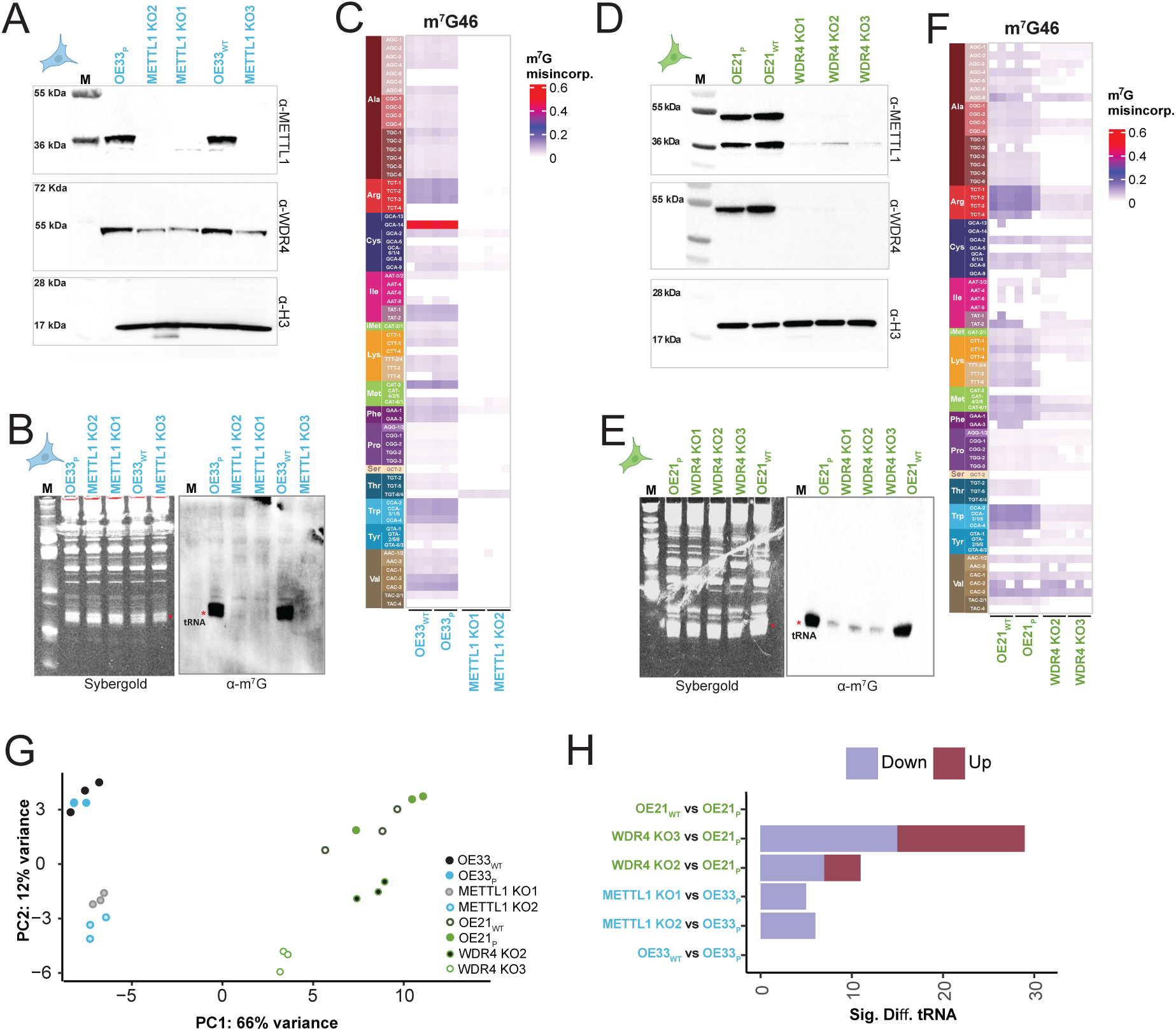
Characterization of METTL1-sensitive OE21 (*WDR4* KO) and METTL1-resistant OE33 (*METTL1* KO) clones A. Immunoblot analysis of parental OE33_P_, Wild-type single cell clone (OE33_WT_) and METTL1 knockout single cell clones (*METTL1* KOs). Whole-cell lysates from ∼75% confluent cultures were transferred to nitrocellulose membrane and sequentially probed for METTL1, WDR4 and H3 (loading control) with specific antibodies. M: MW marker. B. Global m^7^G methylation in OE33 *METTL1* knockout clones. 3 micrograms of total RNA from parental OE33_P_ cells, wild-type clone (OE33_WT_), and individual *METTL1* KOs clones were separated on a TBE–urea gel and stained with SYBR Gold (loading control). RNA was transferred to a nylon membrane and probed with an anti–m⁷G antibody. The position of tRNA is indicated (*). C. tRNA m^7^G46 methylome in OE33 *METTL1* knockout clones. Equal amounts of total RNA from parental OE33 cells, a wild-type single-cell clone, and individual *METTL1* KOs were treated with sodium borohydride followed by aniline cleavage and deep sequencing. Heatmap represents misincorporation at m^7^G46 across tRNAs in the wild-type clone (OE33_WT_), parental line (OE33_P_), and two independent METTL1 knockout clones (METTL1 KO1–KO2). Aminoacids are colour coded (bands at left) and distinct shades indicate different isoacceptors. Isodecoders are identified by their number. Columns correspond to the indicated cell lines. Colour scale denotes relative m^7^G46 signal. D. Immunoblot analysis of parental OE21p, Wild-type single cell clone (OE21_WT_) and WDR4 knocked-out single cell clones (WDR4 KOs). Whole-cell lysates from ∼75% confluent cultures were transferred to nitrocellulose membrane and sequentially probed for WDR4, METTL1 and H3 (loading control) with specific antibodies. M: MW marker. E. Global m^7^G methylation in OE21 WDR4 knockout clones. 3 micrograms of total RNA from parental OE21p cells, a wild-type clone (OE21_WT_), and individual WDR4 knockout clones were separated on a TBE–urea gel and stained with SYBR Gold (loading control). RNA was transferred to a Nylon membrane and probed with an anti–m⁷G antibody. The position of tRNA is indicated (*). F. tRNA m^7^G46 methylome in OE21 *WDR4* knockout clones. As in C. G. Principal component analysis of steady-state tRNA levels (abundance) in OE33 and OE21 derivatives. PCA of normalized tRNA abundance profiles for wild-type OE33_WT_, parental OE33_P_, *METTL1* KO1–KO2, wild-type OE21_WT_, parental OE21_P_, and *WDR4* KO2–KO3. Each point represents an independent replicate. Axes show PC1 and PC2 (PC2 accounts for 12% of the variance); colours correspond to the genotypes indicated in the legend on the right side. H. Global changes in tRNA abundance across OE33 and OE21 derivatives. Stacked bar chart showing the number of tRNA species with significant changes in steady-state levels for the indicated pairwise comparisons among OE33 and OE21 samples described in C and F. Bars are partitioned into tRNAs increased (Up, red) or decreased (Down, purple) in the first genotype listed relative to the second. Significance was defined as an adjusted *P* value < 0.01.

Despite repeated attempts, CRISPR-Cas9 *METTL1* knockout clones in OE21 were not viable. Targeting METTL1’s cofactor, WDR4, resulted in high cell death and, among the surviving clones 87% retained the wild-type *WDR4* allele in heterozygosis (Figure S2C and Methods). Nevertheless, we obtained a few homozygous knockouts (*WDR4*KO), which exhibited varying degrees of proliferative defects, mostly resembling those observed in the parental cell line upon METTL1 siRNA downregulation (Figure S2D). As expected, knockout clones show no sign of WDR4 protein in contrast to the parental line and a wild-type single cell clone (Figure 2D). Notably, loss of WDR4 led to a marked depletion of METTL1 in every *WDR4* KO clone examined (Figure 2D) and a severe reduction in the total amount of m^7^G on tRNAs as assessed by North-Western (Figure 2E). To confirm that those clones are trustworthy METTL1 hypomorphs, we also monitored the methylation on tRNAs at position 46 by mim-tRNAseq profiling, confirming the reduction of m^7^G46 on tRNAs for many although not all isodecoders (Figure 2F).

We then inspected the effect that lack of m^7^G46 may have on the steady state tRNAs level in OE21, OE33 and their corresponding derivatives *METTL1/WDR4* KOs. Northen-blot analysis revealed that deletion of *METTL1* in OE33 did not lead to significant reduction of any of the m^7^G46 modified tRNAs tested (Figure S2E). An oligo probe against another POLIII transcript—5S ribosomal RNA—was used, on the same membrane, as internal control (Figure S2E). Furthermore, no reduction was observed when using probes against a non METTL1/WDR4 substrate such as tRNA-His, which harbours an adenosine at position 46 (Figure S2E). In contrast to OE33 and consistent with previous reports in other cell lines, several m⁷G46-modified tRNAs were reduced in OE21 *WDR4* KO cells, whereas 5S rRNA and tRNA-His remained unchanged (Figure S2F).

Quantification by mim-tRNAseq of steady-state levels of individual tRNAs revealed, as expected, clear cell-line specific expression patterns. The PCA shows the parental and its corresponding WT single-cell clone clustering closely together (Figure 2G, OE21_P_ *vs* OE21_WT_ and OE33 _P_ *vs* OE33_WT_). Given that the wild-type clones (OE21_WT and_ OE33_WT_) have undergone the same selection procedure than KO clones (Figure S2A and S2C), we attribute the following changes in the KO cell-lines to the lack of METTL1/WDR4 activity.

Strikingly, significant differences in tRNA abundance between KO and WT clones encompassed both modified and unmodified tRNAs and occurred irrespective of the METTL1-sensitive or-resistant nature of the cell line (Figure 2H). Yet, at the isoacceptor level, the sensitive OE21 *WDR4* KOs displayed a greater number of affected tRNAs and more pronounced alterations, particularly in the slowest proliferating *WDR4* KO3 (Figure 3A). Consistent with our northern blot data, the most vulnerable isotypes were Val, Lys, and Ala, each showing a significant reduction in their cellular pools despite partial retention of m^7^G46 activity in *WDR4* KOs (Figure 3A). Interestingly, long-term depletion of METTL1/WDR4 activity led to an upregulation of several tRNA isoacceptors—m7G46-and non-m^7^G46 modified—in the OE21 clones. These changes may reflect a response to stress, and an inability to regulate the tRNA pool in the sensitive OE21 cell lines as no upregulation of tRNA pools occurred in the resistant OE33 *METTL1* knockouts (Figure 2H and 3A).

**Figure 3.**
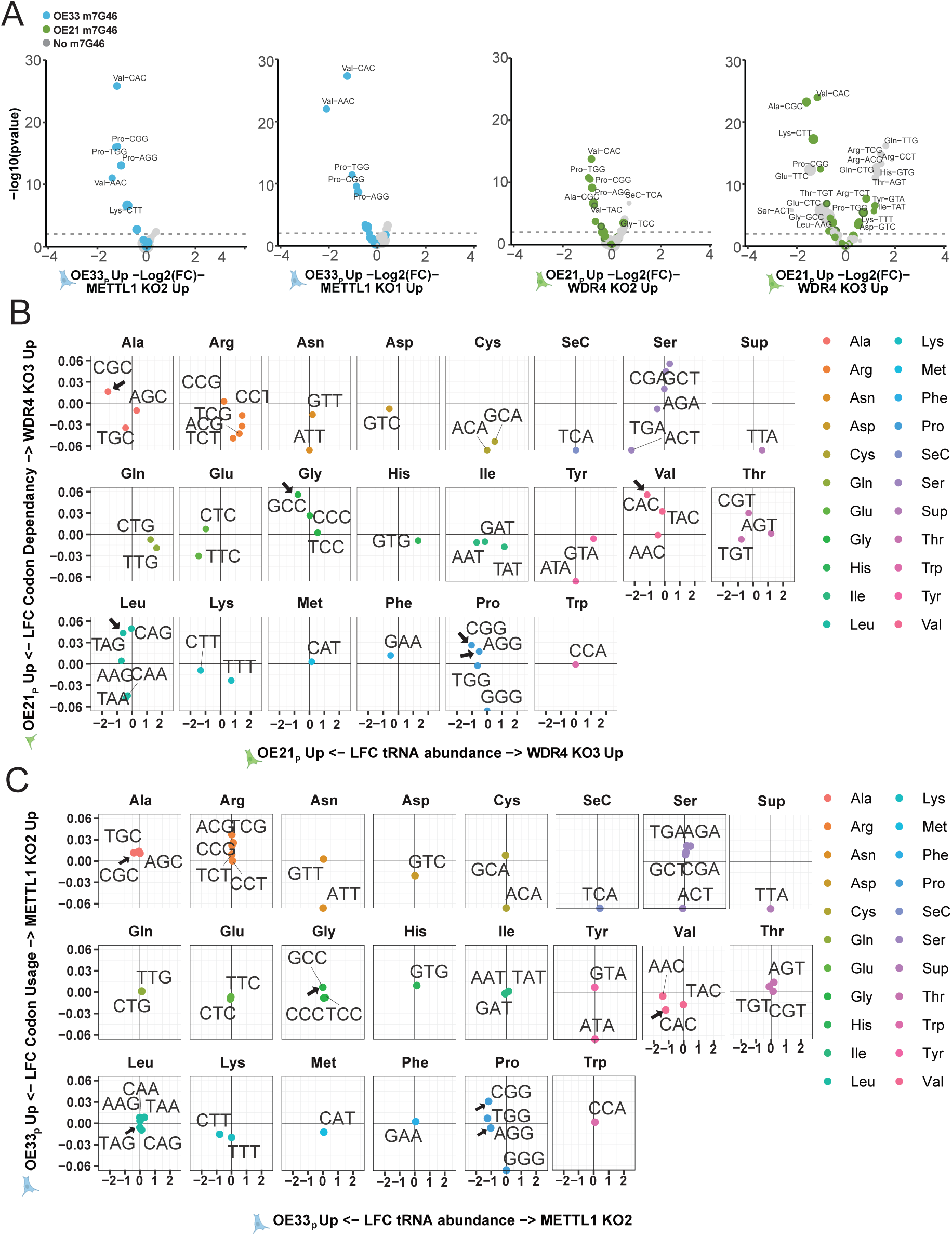
Codon usage *versus* tRNA abundance in OE33 and OE21 derivatives B. Changes in tRNA-isoacceptors abundance across OE33 and OE21 derivatives. Volcano plots comparing parental OE33 (left two panels) or OE21 (right two panels) to their derivatives (knockout clones). Each point is a tRNA isodecoder, coloured by the presence of m^7^G46 (blue, methylated in OE33; green, methylated in OE21; grey, no methylated in any of them). Dot size denote the base mean expression of the issoaceptor family. The x-axis shows log2 fold change (KO vs. parent; positive values indicate higher in the KO, negative values higher in the parent). The y-axis shows −log10(*p*) value; the horizontal dashed line marks the significance threshold (*p* = 0.01). C. Changes in codon demand *versus* cognate tRNA abundance across OE33 and OE21 derivatives. Faceted scatterplots (one panel per amino acid) compare, for each anticodon isoacceptor, the log2 fold-change (KO/parental) in transcriptome-wide codon usage (y-axis) to the log2 fold-change in the abundance of the corresponding tRNA (x-axis). Each point is a single isoacceptor labelled by its anticodon (5′→3′, DNA letters), and coloured by amino-acid identity (key). Vertical and horizontal zero lines denote no change. Points in the upper-left or lower-right quadrants indicate mismatches between demand and supply. Arrows mark isodecoders whose demand is higher that the available cellular pool in the proliferation deficient mutant *WDR4* KO3 (Top panel). The same isodecoders are highlighted in the resistant *METTL1* KO2 (bottom panel). All panels share the same axis scales.

We therefore examined codon usage relative to the abundance of the cognate tRNA isoacceptors in parental OE21 cells and in the slowest-growing *WDR4*KO clone (*WDR4* KO3). Loss of METTL1/WDR4 activity led to a significant increase in the dependency of specific codons decoded by m⁷G46-modified tRNAs—such as Ala-CGC, Val-CAC, Pro-AGG/CGG, Leu-TGA and Gly-GCC. However, the abundance of the cognate tRNAs was markedly reduced in *WDR4*KO3 compared to the parental line (Figure 3B, black arrows). By contrast, OE33 *METTL1*KO2 showed no change in codon usage or abundance of the aforementioned tRNA isoacceptors. When exceptions occurred (e.g., tRNA-Val-CAC), they were parallel at both the tRNA and codon-usage levels (Figure 3C, black arrows). Based on these findings, we propose that the proliferative defect observed in OE21 *WDR4* KO cells may, in part, arise from insufficient supply of specific tRNAs required to meet their translation demands. Whether the changes of non-m⁷G46–modified tRNAs reflects a downstream response aimed at readjusting the cellular tRNA pool to transcriptional changes remains to be determined.

### Lack of tRNA Dihydrouridylation renders OE33 sensitive to METTL1

To gain insights into the genetic network underlying the resistance of OE33 to the lack of METTL1, we performed a genome wide synthetic-lethal CRISPR screen. A wildtype single-cell clone was compared to the two independent *METTL1* knockouts generated with sgRNAs targeting exon 1 or exon 2 (see Figure S2A). Using the Brunello lentiviral library (four sgRNAs per gene, 19,114 protein coding genes)^30^ at a multiplicity of infection of ∼0.2 to assure that each cell is infected with only one virus, we harvested genomic DNA every 4 days over 28 days (Figure S4A and Methods). sgRNAs that were selectively depleted in both *METTL1* KO lines— but not in the wild-type control—were scored as synthetic-lethal hits. Figure 4A shows a Rank plot of the 20000 targeted genes ordered by differential sgRNA dropout (see also Figure S4B). Striking, among the 5 top candidates, we found *POLR3E* (component of the POLIII holoenzyme that transcribe all tRNAs) and the sole nuclear-export receptor for mature tRNAs, *XPOT*, attesting for the specificity of our screen. The other 3 top candidates were in fact tRNA modifying enzymes: The 5-methyl Cytosine, *NSUN2* and two members of the family of Dihydrouridine synthases, DUS-L. A nine-box square interaction plot summarizes the proliferative impact of each hit, with zero on either axis denoting no effect (Figure 4B). Consistent with its role in tRNA export, XPOT loss inflicts a significant growth penalty in the wild-type clone that becomes far more pronounced in the *METTL1*KO background. NSUN2, whose influence on proliferation varies across cancer cell lines, shows a similar context-dependent effect. By contrast, depletion of DUS-L family members only slightly slows the growth of wild-type OE33 cells but markedly impairs proliferation when combined with METTL1 loss (Figure 4B).

**Figure 4.**
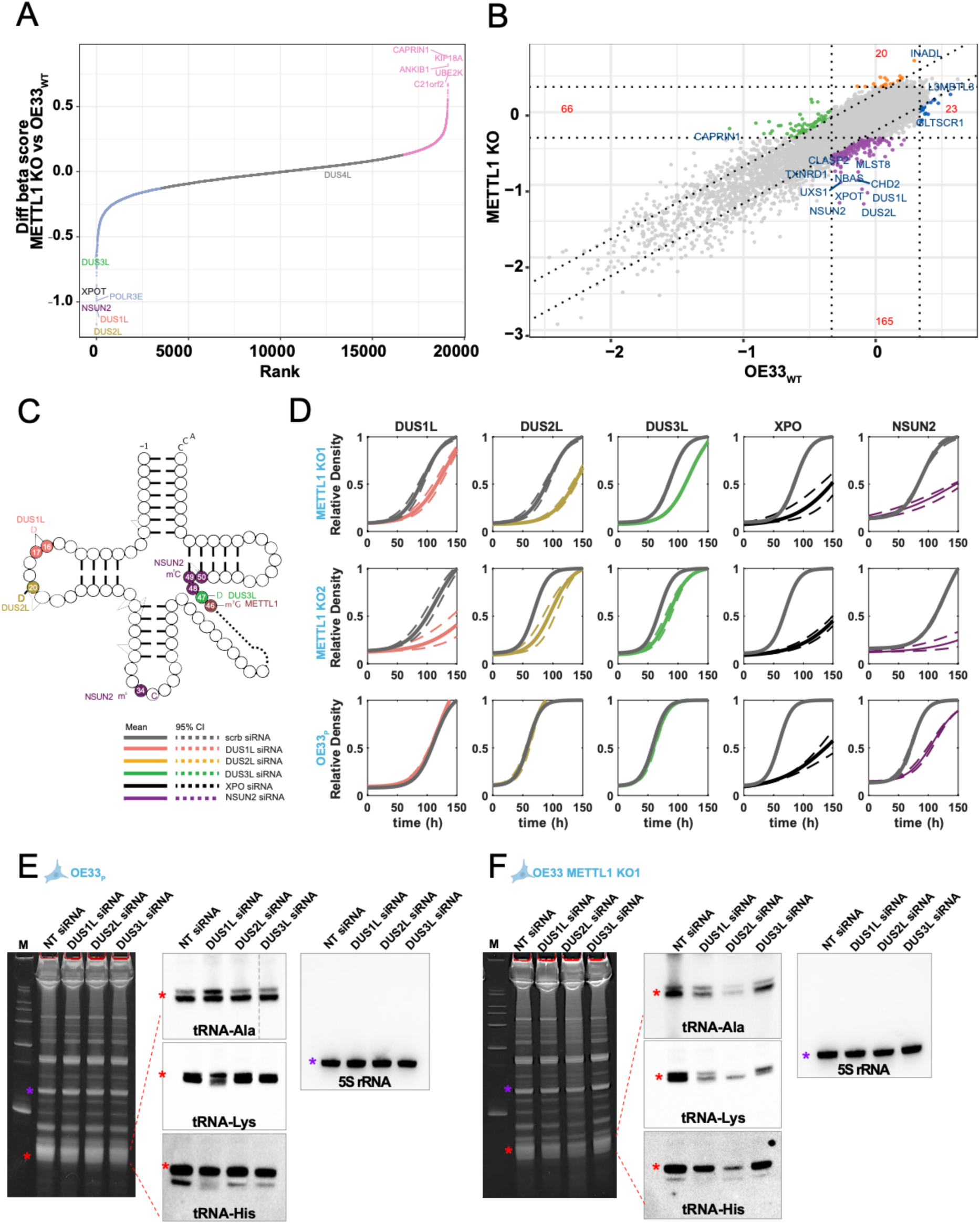
Synthetic lethal screen reveals the Dihydrouridine synthase family as essential in the absence of METTL1/WDR4. A. CRISPR screen ranking for METTL1 dependency. Rank plot of differential β scores (*METTL1* KOs – OE33_WT_) for all targeted genes. Each dot is a gene, ordered from most depleted (left) to most enriched (right). Negative values (dropouts in *METTL1* KOs) are shown in blue; positive values (enrichment in *METTL1* KOs) are shown in pink. Selected top-ranking genes are labelled; label colours denote gene identity as indicated in the colour key. B. CRIPSR 9-square scatter plot displaying the gene selection between OE33_WT_ (x axis) and *METTL1* KOs (y axis). Each axis represents the beta score, and the plot is divided into nine squares, each representing a different combination of positive and negative selection in the two conditions. Negatively selected genes upon *METTL1* KO that appear neutral in the OE33_WT_ appear in the bottom centre square and are shown as purple dots. C. Schematic illustrating the residues modified by the main hits from the synthetic lethal screen in OE33_WT_ and *METTL1* KOs single cell clones. The RNA modifying enzymes are shown in the corresponding colour code. D. Proliferation rates of OE33 *METTL1* KO/ DUS-L siRNA clones. METTL1 KO1 (top), *METTL1* KO2 (middle) and parental OE33_P_ (bottom) cells were transfected with siRNA targeting the main hits from the SL screen (Left to right: DUS1L-siRNA, DUS2L-siRNA, DUS3L-siRNA, XPOTT-siRNA and NSUN2-siRNA) or a non-targeting scramble siRNA as control (grey: scrb. siRNA). Cells were seeded at ∼10% confluency, and growth was monitored every 3 hours using Incucyte. Plots show the proliferative rate (log Δr) normalized to the OE33 wild-type single-cell clone. Dotted lines represent 95% CI. (n=3). E. Northern blot analysis of tRNA from parental OE33_P_ and OE33_P_ DUS-L knockdown clones. OE33p was transfected with 2.5nM siRNA targeting DUS1L-siRNA, DUS2L-siRNA, DUS3L-siRNA or a non-targeting scramble siRNA as control (scrb. siRNA). Cells were collected 5 days post-transfection. From each sample, 3 micrograms total RNA were resolved in TBE-Urea gels stained with Sybergold (left panels), transferred to Nylon membranes and hybridized, firstly with oligos against tRNAs (as indicated) and subsequently with an oligo against the POLIII transcript ribosomal 5S, used as additional loading control. tRNA are marked by a red asterisk (**\***), r5S is marked by a purple asterisk (**\***) F. Northern blot analysis of tRNA from *METTL1* KO1 and *METTL1* KO1 DUS-L knockdown clones. *METTL1* KO1 was transfected with 2.5nM siRNA targeting DUS1L-siRNA, DUS2L-siRNA, DUS3L-siRNA or a non-targeting scramble siRNA as control (scrb. siRNA). Cells were collected 5 days post-transfection. From each sample, 3 micrograms total RNA were resolved in TBE-Urea gels stained with Sybergold (left panels), transferred to Nylon membranes and hybridized, firstly with oligos against tRNAs (as indicated) and subsequently with an oligo against the POLIII transcript ribosomal 5S, used as additional loading control. tRNA are marked by a red asterisk (**\***), r5S is marked by a purple asterisk (**\***)

In terms of gene ontology, the analysis revealed a strong correlation to RNA biology with tRNA dihydrouridine synthesis and tRNA modifications as top gene networks related to METTL1 function (Figure S4C).

Remarkably, out of more than 40 modifications described on human tRNAs, dihydrouridylation at all canonical positions on the “D loop” (D16, D17, D20) and m^5^C (which decorates the anticodon at position 34 and the junction between the variable and “T loop”) displayed the strongest functional redundancy with m^7^G46. These findings suggest that a coordinated tRNA structural compensation mechanism enables OE33 cells to withstand METTL1 loss (Figure 4C).

We validated the results of our CRISPR-Cas9 SL screen by efficient siRNA downregulation of our hits (Figure S4D). In agreement with the above results, siRNA of XPOT and NSUN2 resulted in obvious proliferative defect in the parental OE33, which became more severe when combined with *METTL1* KO (Figure 4D). Since parental OE33 cells are highly dependent on NSUN2—and because NSUN2 deposits m^5^C at numerous tRNA positions^31^—we desisted to further investigate the molecular interplay between METTL1 and NSUN2 on tRNA in our model system.

Instead, we focused on the dihydrouridine synthase (DUS-L) family, which comprises four evolutionarily conserved enzymes with strict positional specificity on tRNA substrates: DUS1L reduces U16/17 in the D-loop, DUS2L modifies U20, DUS3L acts on U47 at the variable loop, and DUS4L targets the non-canonical U20a/U20b (Figure 4C)^32^. DUS1L and DUS2L ranked among the five strongest synthetic lethal hits, and DUS3L also scored prominently (Figure 4A). We therefore validated these three hits: while DUS1L, DUS2L, and DUS3L downregulation caused negligible growth delay in parental OE33 cells, it strongly impaired proliferation in the *METTL1* KO background, confirming the CRISPR-based synthetic lethal interaction (Figure 4D). Concurring with the proliferation, cellular levels of m^7^G46 modified tRNAs —tRNA-Lys and tRNA-Ala —were unaffected by down regulation of the DUSL enzymes in the parental OE33, except a mild decrease upon DUS1L (Figure 4E). In contrast, these tRNAs were markedly reduced in OE33 *METTL1*KO cells treated with DUS-L siRNA (Figure 4F), resembling the effect of METTL1/WDR4 loss in the sensitive OE21 line (Figure S2F). Noteworthy, a non METTL1 substrate (tRNA-His) was only marginally affected by the combined METTL1/DUS-L reductions, except upon DUS2L siRNA likely due to the extremely severe growth defect (Figure 3F). Hence, we confirmed the synthetic lethal relationships disclosed by our CRISPR-Cas9 screen and conclude that absence of dihydrouridilation renders the naturally resistant OE33, now sensitive to lack of METTL1.

### A balance between m7G46 and dihydrouridine defines optimal tRNA structure

To assess how the activities of METTL1/WDR4 (m⁷G46 methyltransferase) and the DUS-L (dihydrouridine synthases) influence tRNA architecture, we would ideally investigate the structure of individual tRNAs isolated from METTL1KO clones down-regulated for individual and combined DUS-L enzymes. However, the impact on viability of such combinations made impossible to generate enough RNA to attempt structural work. Thus, we examined a synthetic oligo corresponding to tRNA-Tyr-GTA-1. This tRNA is known to be modified at D16, D17 & D20 (classical D-loop positions), harbours m⁷G46 directly 5′ to D47 at the variable loop/hinge and contains a UUCGAA tetraloop-like motif rather than the canonical TΨC/m¹A58 triad at the T-arm/loop. Oligos corresponding to this mature tRNA were synthesized in 2 forms: (i) unmodified and (ii) containing D residues at positions 16,17,20 and 47 plus m⁷G46 (D16D17D20m^7^G46D47) (Figure S5A). Far-UV circular dichroism (CD) spectra confirmed that both RNAs adopt A-form helical characteristics typical of folded tRNAs (Figure 5A). CD melting curves at 270 nm showed single, cooperative transitions for both RNAs (Figure 5B and Figure S5B). Using the half-signal criterion ^33^, the D16D17D20m^7^G46D47 modified tRNA exhibited the highest Tm (61.6 °C), exceeding the unmodified (57.4 °C). In addition, D16D17D20m^7^G46D47 markedly sharpened the melt implying enhanced cooperativity, consistent with reinforcement of tertiary contacts (Figure 5B). These data suggest that the presence of the above modifications strengthens tertiary contacts at the elbow. The resulting balance between rigidity (cooperativity/thermal-resilience) provided by m^7^G46 and plasticity (breathable folding) contributed by Ds, likely tunes downstream steps—including aminoacylation, deposition of additional modifications, EFTu binding, and ribosome accommodation— the disruption of which may underlie the lethality observed upon simultaneous abrogation of METTL1/WDR4 and DUS-L activities.

**Figure 5.**
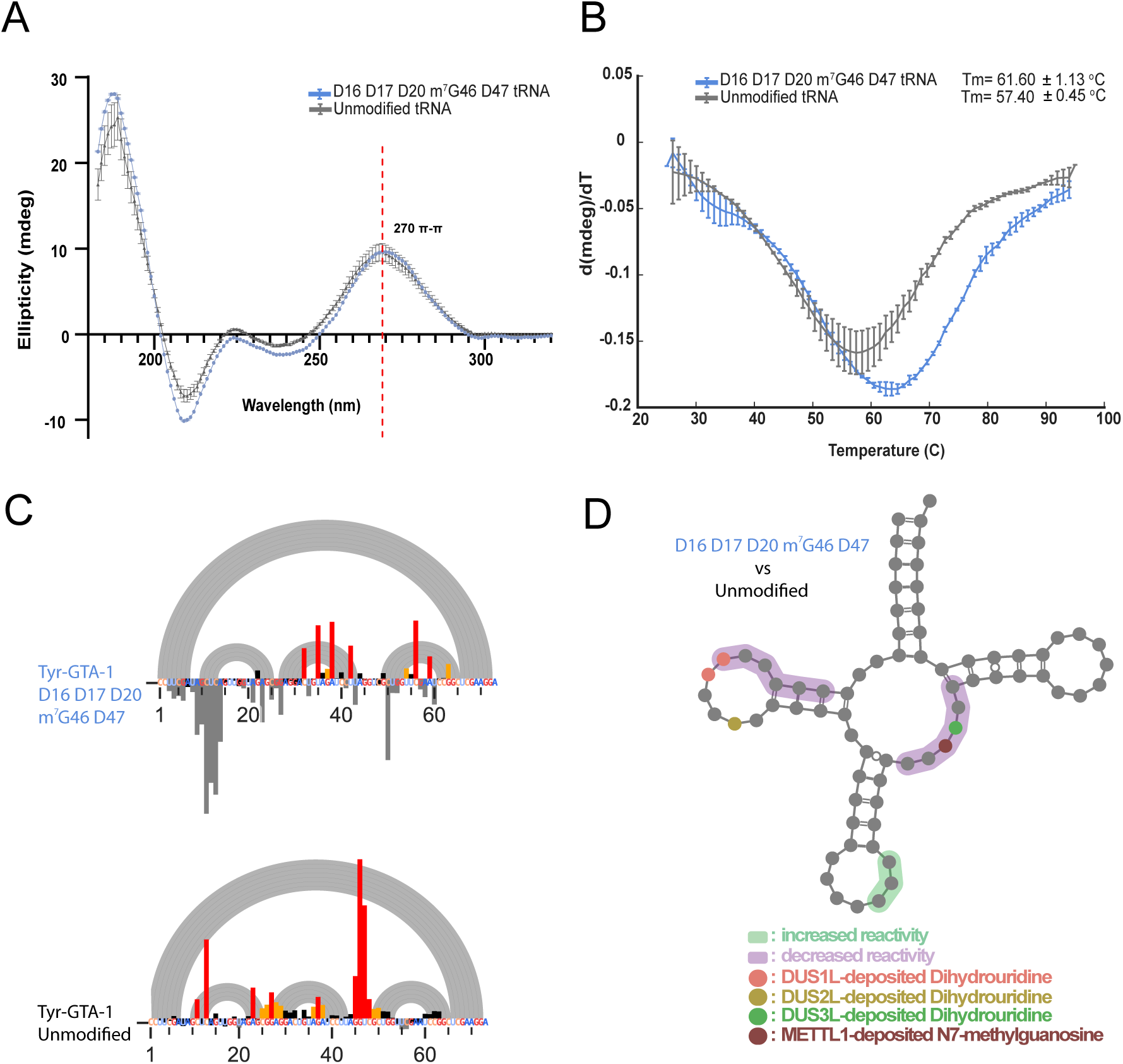
Dihydrouridine and m^7^G46 cooperate to provide optimal tRNA structure and stability **A.** Circular Dichroism (CD) Spectra of Modified and Unmodified Tyr-GTA-tRNA. Spectra are shown for tRNA containing D16D17D20m^7^G46D47 (blue) or unmodified tRNA (grey). Measurements were performed at 25 °C in sodium phosphate buffer, with ellipticity plotted as a function of wavelength (nm). **B.** Tyr-GTA-tRNA Thermal Denaturation Profiles. First derivatives of normalized CD melting curves at 270 nm reveal the apparent melting temperatures (Tm) for tRNA oligos carrying distinct modification patterns. Data are shown for tRNA containing D16D17D20m^7^G46D47 (blue) or unmodified tRNA (grey). Peaks in the negative derivative correspond to the primary cooperative unfolding transitions. Data are representative of 2 independent replicates. **C.** SHAPE-MaP probing of synthetic tRNA-Tyr variants. Arc plots (grey) depict canonical base-pairing interactions. Bar graphs (black to red bars) show per-nucleotide SHAPE reactivity for the dihydrouridine plus m^7^G46 tRNA (D16D17D20m^7^G46D47), and the unmodified one with a colour scale from black (low reactivity) to red (high reactivity). **D.** Schematic secondary structures of tRNA-Tyr highlight the positions of dihydrouridines and m^7^G46, with nucleotides shaded according to observed reactivity changes (purple: protections, green: enhancements).

To further investigate the effect of the m^7^G46 and dihydrouridine on tRNA structure at the nucleotide resolution, we used 1-Methyl-7-nitroisatonic anhydride (1M7) chemical probing coupled to mutational profiling (SHAPE-MaP)^35^, a widely used method for RNA secondary structure analysis. Chemical probing has been previously applied to tRNAs to study *in vitro* and in cell^34–36^. Since the lack of m^7^G46 plus dihydrouridine results in a substantial reduction of tRNAs in the cellular pool, it was not technically feasible to probe their structure on cell-isolated material. Instead, we relied on the same synthetic tRNAs that were used for CD analysis.

In SHAPE-MaP, low reactivity indicates base-paired or conformationally constrained nucleotides, whereas high reactivity marks flexible, solvent-exposed positions. In agreement with the thermal data, the D16D17D20m^7^G46D47 oligo shows the most native-like pattern: uniformly low reactivity across helices, pronounced protection near 46 (variable/T region) and within the D/T-loop core, consistent with stabilization of the elbow via the C13–G22–m⁷G46 base triple and long-range D–T contacts; loop signals remaining high (Fig. 5C, top). In contrast, the unmodified RNA exhibits elevated reactivity across the D stem and variable loop—indicating secondary structure instability and poor tertiary packing (Fig. 5C, bottom).

ΔSHAPE (modified − unmodified) analysis highlights protection of the D-stem/loop and variable loop (purple shading), consistent with variable loop docking towards the D-arm upon m⁷G46 installation, and optimal anticodon-loop exposure (green shading) in the fully modified RNA. Simultaneous lack of Ds and m^7^G46 results in unstable secondary structure and poor tertiary folding (D/T loop core not formed and suboptimal anticodon exposure), underscoring the observed effect that simultaneous lack of METTL1/WDR4 and dihydrouridine synthases has on cell fitness.

### Crosstalk between DUS3L and METTL1 buffers loss of tRNA m⁷G46

In addition to N7-methyl guanosine, mim-tRNAseq detects other RNA nucleoside modifications such as m¹A, m¹G, m^2,2^G and dihydrouridine. Our mim-tRNAseq dataset revealed that loss of METTL1 in the resistant OE33 line upregulates DUS3L activity at its tRNA substrates. The increase in D at position 47 occurs in a substantial pool of m^7^G46 modified tRNAs isoacceptors, in both OE33 *METTL1*KOs respect to the wild-type clone and parental cell line (Figure 6A, blue label). Specifically, several isoacceptor families for Val, Lys, Tyr, Pro, Cys, Phe, Met, Trp and iMet exhibit increased D47 in OE33 upon *METTL1* deletion. The effect is notorious across the m^7^G46 modified tRNA pool, even at the isotype level and it does not occur on non-m^7^G46 modified tRNAs suggesting a response to the lack of METTL1 activity in the resistant OE33 cell line (Figure 6B, OE33).

**Figure 6.**
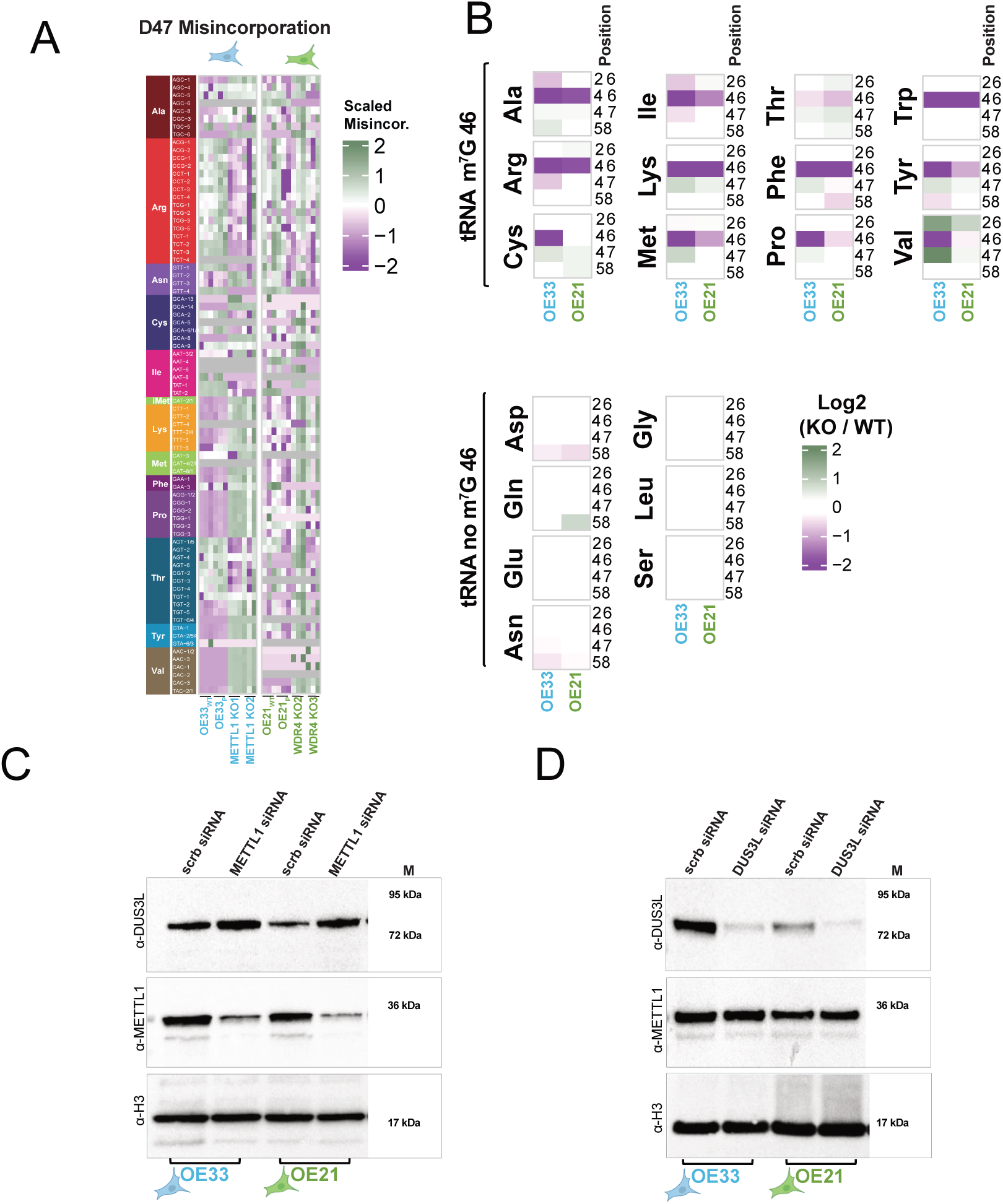
Lack of METTL1/WDR4 results in increased DUS3L protein and D47 on m^7^G46 modified tRNAs. A. Heatmap of dihydrouridine at position 47 (D47) on tRNAs of resistant OE33_P_ and sensitive OE21_P_ parental cell lines and derivatives. Changes in D47 are calculated from reverse-transcription misincorporation rates after sodium borohydride (NaBH₄) treatment and deep sequencing (tRNA mim-seq). tRNA isotypes and isodecoders are colour coded and listed on the left side of the maps. In blue OE33_P,_ wild-type OE33_WT_, *METTL1*KO1 and *METTL1*KO2; In green parental OE21_P_, wild-type OE21_WT_, *WDR4*KO2 plus *WDR4*KO3. B. tRNA-isotype modification changes in *METTL1* KO OE33 (R) and *WDR4* KO OE21 (S). Heatmap of isotype-level changes in m2,2G26, m^7^G46, D47, and m1A58 quantified by tRNA mim-seq. Values are KO/reference ratios, where the reference is the combined parental + wild-type clone for each line (OE33, blue; OE21, green). Top: m^7^G46-modified isotypes. Bottom: non-m^7^G46 isotypes. C. Immunoblot analysis of DUS3L response to METTL1 knockdown. OE33 and OE21 parental cell lines were transfected with siRNA targeting METTL1 or non-targeting scramble siRNA (scrb. siRNA). Cells were collected 5 days post-transfection, and 45 micrograms of total protein extract were transferred to Nitrocellulose membrane and immunoblotted firstly with anti-DUS3L and sequentially with anti-METTL1 and anti-Histone H3 as a loading control. D. Immunoblot analysis of METTL1 response to DUS3L knockdown. OE33 and OE21 parental cell lines were transfected with siRNA targeting DUS3L or non-targeting scramble siRNA (scrb. siRNA). Cells were collected 5 days post-transfection, and 45 micrograms of total protein extract were transferred to Nitrocellulose membrane and immunoblotted firstly with anti-DUS3L and sequentially with anti-METTL1 and anti-Histone H3 as a loading control.

Alongside elevated D47, the resistant knockout clones displayed altered stoichiometry of m²²G26, m¹A58, and D20 (Fig.6B and Figure S6A). These changes though, are restricted to scarce members within few isodecoder families. Precisely, *METTL1*KO resistant clones show an increase in m^1^A58 and decreased m^2,2^G26 on tRNAs for Alanine—one of the few isotypes less responsive *via* increasing D47—suggesting a distinct sequence-dependent strategy for tRNA stabilization. In clear contrast to the OE33 *METTL1* KOs, no overall patters are observed in OE21 *WDR4* KOs regarding the mentioned modifications, apart from the reduction in m^7^G46, (Figure 6A, 6B and Figure S6A). Thus, the METTL1 resistant, but not the sensitive cell line, responds to lack of METTL1/WDR4 activity by re-wiring the tRNA epitranscriptome mainly by increasing the D47 tRNA content. Consistent with the above, the responsible enzyme DUS3L is upregulated at the mRNA and protein levels in OE33 but not OE21 knockouts (Figure S6B, S6C and S6D). These findings suggest that DUS3L upregulation may contribute to the intrinsic METTL1 resistance of OE33 cells.

To explore whether the crosstalk between METTL1 and DUS3L was a fast cellular response, we performed METTL1 siRNA downregulation in the parental OE33 line and assessed DUS3L expression 4 days post-transfection. Western blotting confirmed robust METTL1 depletion and a concomitant DUS3L increase, mirroring the effect observed in OE33 *METTL1*KO clones (Figure 6C).

Unexpectedly, the METTL1-sensitive OE21 line also upregulated DUS3L upon METTL1 knockdown, unlike OE21 *WDR4*KO clones, which were non-responsive (Figure 6C and Figure S6D).

Interestingly, the METTL1–DUS3L crosstalk does not work reciprocally: siRNA mediated DUS3L knockdown does not change the METTL1 protein level, neither in the resistant nor in the sensitive cell line (Figure 6D). Thus, whereas loss of DUS3L (and consequently D47) does not alter METTL1 levels, METTL1 depletion induces DUS3L upregulation, consistent with a compensatory response to the absence of m^7^G46 on tRNAs.

### METTL1 resistant cell lines express high DUS3L protein levels

DUS3L upregulation upon METTL1 siRNA in OE21 was intriguing given the lack of response in the *WDR4*KO clones which, otherwise, phenocopied the former one. However, we noticed that the endogenous DUS3L protein level was substantially different in OE33 and OE21 cell lines (Figure 6C and 6D, scramble siRNA, lines 1 and 3). That was confirmed by interrogating our RNA-seq dataset (Figure S7A). To rule out subtle metabolic differences, we prepared extracts from parental OE21 and OE33 cells grown to ∼50% or 100% confluence (both lines have the same medium requirements). We found that regardless confluency, DUS3L levels were reproducibly higher in OE33 than in OE21 (Figure 7A). To test whether low DUS3L was a singularity of OE21 or there was a correlation between its expression and METTL1 resistance, we interrogated our panel of cancer cell lines showing different dependency on METTL1 for proliferation (see Figure S1). We observed that those cell lines resistant to METTL1 down-regulation exhibited higher DUS3L protein levels than the sensitive ones. In fact, DepMap database revealed a significant correlation between METTL1 sensitivity (by CRISPR-Chronos score) and DUS3L expression in our cell panel (Figure S7B). Altogether, the resistance to METTL1 depletion seems to correlate (to the extent of our cell panel) to high DUS3L protein levels.

**Figure 7.**
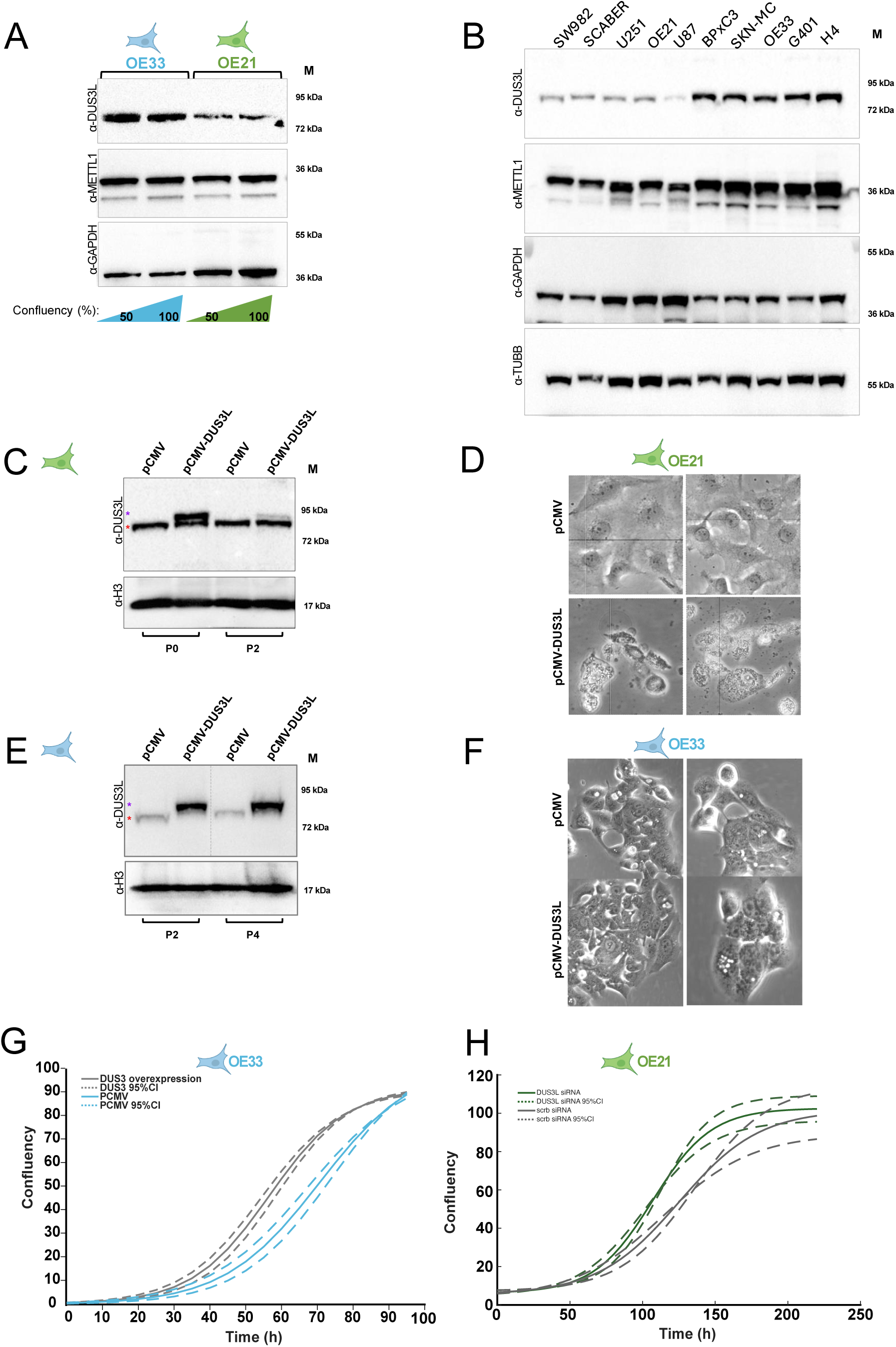
METTL1 resistance correlates with high DUS3L protein expression. B. Immunoblot analysis of DUS3L levels in OE33_P_ and OE21_P_ parental cell lines. Total protein extracts (45 micrograms) from cells grown at 50% and 100% confluency, were transferred to Nitrocellulose membrane and immunoblotted firstly with anti-DUS3L and sequentially with anti-METTL1 and anti-GAPDH as a loading control. C. Immunoblot analysis of DUS3L levels across the panel of Cancer Cell Lines shown in Figure 1A. Total protein extracts (45 micrograms) from cells grown at 75% confluency, were transferred to Nitrocellulose membrane and immunoblotted firstly with anti-DUS3L and sequentially with anti-METTL1 and anti-GAPDH as a loading control. D. Immunoblot analysis of endogenous DUS3L and overexpressed myc-DUS3L in parental OE21. Total protein extracts (45 micrograms) from stable multiclonal population generated by transfection of empty-pCMV vector or pCMV-myc-DUS3L plasmids, were transferred to Nitrocellulose membrane and immunoblotted firstly with anti-DUS3L (note signal acquired after 37 sec exposure) and sequentially anti-H3 as a loading control. Endogenous DUS3L is marked with red asterisk (**\***), transfected myc-DUS3L is marked with purple asterisk (**\***) E. Representative light-microscopy images of OE21 cells with endogenous DUS3L (empty-vector control, pCMV; top) or DUS3L overexpression (pCMV-myc-DUS3L; bottom). Images were acquired 4 days after completion of antibiotic selection of transfected cells F. Immunoblot analysis of endogenous and pCMV-overexpressed DUS3L in parental OE33. Total protein extracts (45 micrograms) from stable clones generated by transfection of empty-pCMV vector or pCMV-myc-DUS3L overexpression plasmid cells, were transferred to Nitrocellulose membrane and immunoblotted firstly with anti-DUS3L (note signal acquired after 5 sec exposure) and sequentially anti-H3 as a loading control. Endogenous DUS3L is marked with red asterisk (**\***), transfected myc-DUS3L is marked with purple asterisk (**\***) G. Representative light-microscopy images of OE33 cells with endogenous DUS3L (empty-vector control, pCMV; top) or DUS3L overexpression (pCMV-myc-DUS3L; bottom). Images were acquired 4 days after completion of antibiotic selection of transfected cells. H. Proliferation rates of parental OE33 stable clones expressing either endogenous (empty-pCMV, grey) overexpressed DUS3L levels (pCMV-myc-DUS3L, blue), as in panel E. Cells were seeded at ∼10% confluency, and growth was monitored every 3 hours using Incucyte. (n=6) I. Proliferation rates of parental OE21 cells transfected with either scramble control siRNA (grey) or DUS3L siRNA (green). Cells were seeded at ∼10% confluency, and growth was monitored every 3 hours using Incucyte. (n=3)

### The METTL1 sensitive OE21 cell line cannot sustain high DUS3 expression

A prediction from the above would be that DUS3L overexpression could convert OE21 into METTL1 resistant. To test this hypothesis, we created a OE21 stable multiclonal population by integrating either a vector driving constitutive overexpression of DUS3L under the strong CMV promoter, or the control “empty vector”. Antibiotic selection inflicted severe cell death in the case of pCMV-DUS3L. Among the survivals, as expected, we detected the transfected DUS3L allele — which runs higher due to a myc-tag— only in the relevant population, whereas the endogenous protein showed up in mock and DUS3L overexpressing clones (Figure 7C). However, after 2 cell passages (12 days), the pCMV-DUS3L transfected population shows barely any sign of myc-DUS3L regardless of their resistance to the antibiotic used for selection. Thus, likely, the surviving cells have simply repressed the expression of DUS3L from the pCMV-promoter. In fact, five days after transfection, the mock (pCMV-empty) showed no loss of fitness, whereas pCMV-DUS3L expression caused clear toxicity and reduced cell health (Figure 7D). OE21’s inability to maintain high DUS3L protein levels may explain the discrepancy between the transient response to METTL1 siRNA and the prolonged deprivation in *WDR4*KOs.

Remarkably, expression of pCMV-myc-DUS3L drove sustained robust DUS3L overexpression in OE33 cells—far exceeding its endogenous levels (Figure 7E; compare endogenous and overexpressed proteins)—without inducing cell death (Figure 7F). Moreover, DUS3L overexpression significantly increased the proliferation rate of OE33, indicating that tolerance to high DUS3L levels contributes to cellular fitness. Conversely, OE21 showed a similar proliferative advantage, but only after siRNA-mediated DUS3L knockdown (Figure 7G). Collectively, these data suggest that METTL1 sensitivity is driven, at least in part, by the inability of certain cell lines to sustain elevated DUS3L protein levels.

In summary, our findings reveal that OE21 vulnerability to METTL1 loss arises from the combination of its dependency on cell-line specific m⁷G46-modified tRNAs and the restricted capacity to rewire the tRNA epitranscriptome, particularly D47, to compensate for lack of m⁷G46.

## DISCUSSION

### METTL1/WDR4 governs tRNA stability and translational programs in cancer cells

In mammals, METTL1 partners with WDR4 to install m^7^G46 across a defined subset of tRNAs, stabilizing their cellular pool and reprograming translation in favour of codons read by m^7^G46-modified tRNAs—often enriched in growth and stress-response genes^13,14,37^. Hence, overexpression of METTL1/WDR4 is common across many different types of cancers, making this complex an appealing therapeutic target^17–21^. METTL1/WDR4 dosage and enzymatic output are rate-limiting: small changes in activity can push target tRNAs past a stability threshold, triggering pronounced phenotypes without globally collapsing translation^17,38^. Yet, METTL1/WDR4 expression and dependency do not always track linearly across tumour types, and phenotypes vary with codon usage, tRNA isodecoder expression, and cooperating tRNA modifications^39,40^. We identified a panel of cell lines that exhibit a very variable range of dependency on METTL1/WDR4. The impact of METTL1 perturbation is highly context dependent: striking vulnerabilities emerged in glioblastoma (U251, U87), Synovial Sarcoma (SW982) and squamous carcinoma (OE21) models, where METTL1 disruption markedly reduced proliferative capacity across multiple parameters, whereas kidney (G401), oesophageal adenocarcinoma (OE33), and pancreatic adenocarcinoma (BxPC3) were largely resistant, suggesting that the oncogenic role of METTL1 is not universal, but rather shaped by lineage-specific dependencies on tRNA modification programs.

Among the above, we selected sensitive and resistant oesophagus cell lines (OE21 and OE33) to investigate the molecular bases of METTL1 independency. Our results show small differences between these two cell lines regarding overall tRNA loci transcription and m^7^G46 tRNA methylome. The very few differences in the m^7^G46-methylome between cell lines often reflects their loci usage (eg. tRNA-Arg-TCT-4 or tRNA-Met-CAT-4, are modified only in OE21 because they are expressed only in OE21). The impact of these m^7^G46 modified tRNAs depends on their use to implement each cell line specific transcriptome. For example, there is a significant bias in the use of Ala-AGC or Arg-TCT in OE21 (S), whereas Gln-GTC and Cys-GCA are more prevalent in OE33 (R). Overall, we find a significant bias toward modified codon usage in the sensitive cell line, although both lines ultimately converged in terms of isoacceptor and isotype representation. Furthermore, the abundance of m^7^G46-modified and no modified isoacceptors does not mirror the observed codon bias, implying that few cell-line-specific isodecoders—rather than the bulk— may determine the METTL1/WDR4 vulnerability observed in OE21 and reported in other cell lines^23,38,41^.

Remarkably, poor growing METTL1-hypomorphs (*WDR4* KOs) in the sensitive OE21 line showed increased usage of certain m⁷G46-decoded codons but reduced abundance of the cognate isoaceptors relative to parental OE21. Based on this, we propose that OE21’s sensitivity to METTL1 loss arises, at least in part, from its inability to adjust the cellular tRNA pool to the transcriptional and translational shifts triggered by insufficient METTL1/WDR4 activity. In contrast, OE33 METTL1 knockouts exhibited no codon-usage bias and cognate abundance or there is coordinated decrease on codon-usage and cognate abundance (e.g., decrease in tRNA-Val-GUG codon usage and tRNA-Val-GAC isoacceptor). This suggests compensatory mechanisms operating in this resistant cell line.

To investigate the molecular pathways operating under chronic m⁷G46 deficiency in the resistant line OE33, we analyzed *METTL1* single-cell–knockouts derivatives. We confirmed that no alternative methyltransferase compensates for METTL1 activity, as tRNAs remained devoid of m^7^G46. Despite this, isodecoders destabilized in METTL1-sensitive contexts (like OE21 and other published cell lines) remained mainly unaffected in OE33, consistent with the lack of proliferation defects in *METL1*KO clones.

Our synthetic lethal (SL) screen in the resistant OE33, designed to unmask functional dependencies, uncovered an unexpectedly defined functional axis: of the >40 distinct tRNA modifications known in humans (averaging ∼13 per tRNA)^34^ only 5-methylcytosine (m^5^C) and dihydrouridine (D) emerged as essential for OE33 in the absence of m^7^G46. Although this network points to tRNA structural readjustments, we cannot exclude additional metabolic rewiring or changes in translational control buffering against m^7^G46 loss.

### METTL1/WDR4 as a central node in the tRNA structural network

A conserved structural feature of tRNAs is the 13–22–46 base triple and the D/T-loop “elbow” contacts (e.g., G18–Ψ55 and G19–C56) that stabilize long-range tertiary architecture, defining the L-shaped fold. N7-methylation at G46 introduces a positive charge that increases local hydrophobicity, enhancing electrostatic interactions at the elbow and coaxial stacking of the acceptor and T stems^42^. Consequently, m⁷G46 preserves a native geometry recognized by cognate aminoacyl-tRNA synthetases and protects the 5′ leader from surveillance mechanism that monitor acceptor/T-arm geometry^6,9,11^. METTL1/WDR4 mutations (*TRM8/TRM82* in yeasts) destabilize only a subset of tRNAs, on the bases of (i) identity elements in the acceptor and T stems, (ii) sequence/structural features near positions 13, 22, and 46, and (iii) the local modification landscape that can buffer or exacerbate loss of m⁷G46^6–13,42–46^. Consistent with the above, our synthetic lethal (SL) screen revealed NSUN2 (m^5^C) and three of the four dihydrouridine synthase (DUS-L) enzymes as essential for viability in the absence of m^7^G46.

NSUN2 dependent methylation of 5 cytosine on tRNAs at positions C34 and C48–C50 have been shown important in the maintenance of tRNA structural integrity and fitness in several cell lines^46,47^, as we show in this study for OE33. Yet, concurrent depletion of m⁵C and m⁷G46 precipitates a sharp loss of viability possibly due several non-exclusive effects: **(i)** removal of nearby m⁵C (C48–C50) further erodes local stacking/geometry, dropping the functional tRNA pool below the stability threshold that triggers rapid tRNA decay (RTD) and **(ii)** loss of m⁵C protection against angiogenin-mediated cleavage, causing accumulation of tRNA fragments that repress translation initiation and amplify stress responses^46–48^.

In contrast to NSUN2, the effect of insufficient DUS-L activity is relatively mild. By saturating the C5–C6 double bond of uridine, D increases ring puckering and local backbone flexibility. When placed at positions D16/17 (DUS1L) and D20 (DUS2L), these “soft spots” allow the D-loop to adapt and engage correctly with the T-loop and the variable region, stabilizing the long-range interactions^49,50^. Our SL screen disclosed DUS1L and DUS2L among the 5 top hits. If D16/D17, or D20 are absent the D-loop is expected to become overly rigid, less able to accommodate structural strain when the elbow is weakened (as in m⁷G46 deficiency) leading to misfolding, as shown in our structural study using synthetic Tyr-tRNA oligos. The resulting changes in exposure/occlusion at several helices and stems can explain the severe proliferative phenotype observed in *METTL1* KOs when combined with DUS-L CRISPR-Cas9 knockout (SL screen) or DUS-L siRNA knockdowns.

### DUS3L: An integrated model of modification crosstalk

The third member of the DUS-L family, DUS3L, modifies the uridine adjacent to G46 and scored prominently in our synthetic-lethality screen. As observed for the other dihydrouridine synthases, combined depletion of DUS3L and METTL1 produced a clear synergistic growth defect. In addition, our mim-tRNAseq analysis revealed that OE33 METTL1 knockout cells compensate for the absence of m⁷G46 by upregulating DUS3L transcripts, protein, and D47 modification across a broad number of tRNAs. Many isodecoders for Lys, Met, iMet, Pro, Ile, Thr, Tyr, and Val, exhibited increased D47 content in METTL1-deficient clones compared with parental and wild-type controls. Strikingly, this adaptive response is restricted to the METTL1-resistant OE33 cell line, suggesting a compensatory mechanism that stabilizes its tRNA pool and does not operate in the sensitive OE21 line. Indeed, virtually all OE33 m⁷G46-modified tRNA isotypes displayed increased D47 in the absence of m⁷G46, with the notable exception of alanine. Even in this case, specific tRNA-Ala isodecoders (such as Ala-AGC-1 and Ala-TGC-6) that make a substantial contribution to the OE33 tRNA pool, showed a trend toward increased D47 levels in the METTL1 KOs.

Conceptually, tumours may exploit increased D47 to **(i)** accelerate folding of abundant tRNAs under proteostatic pressure and **(ii)** selectively stabilize tRNAs required for oncogenic codon bias. Structurally, D47—positioned adjacent to G46 in the elbow—introduces local flexibility that enables the D/T-loop to absorb conformational strain and provides additional adaptability at the elbow, thereby compensating for the weakened 13–22–46 base triple and D/T-loop stacking interactions that arise when m⁷G46 is absent^49–51^.

Remarkably, the few isotypes that did not follow the “D47 rule” (some Ala, Cys and Ile isodecoders) use alternative *compensatory elbow stabilizers* such as D20 reduction to increase D-loop stiffness, and —in case of Ala-tRNA isodecoders— increase m1A58 which reinforces T54–Ψ55–A58 interactions to rescue acceptor-stem breathing and aminoacylation defects own to lack of m^7^G46. We also detected changes in m²,²G26, a hinge “lock” between the D-and anticodon stems: it increased in D47-regulated Val isodecoders but decreased in m¹A58-buffered tRNAs, suggesting isoacceptor-specific tuning of tRNA rigidity and flexibility. Notably, these responses are specific to METTL1 loss—the wild-type single-cell clone is virtually identical to parental OE33—and occur exclusively in m⁷G46-modified tRNAs. Strikingly, in the METTL1-sensitive cell line OE21, tRNAs do not change their modification landscape when m^7^G46 decreases. Because loss of METTL1 in this background correlates with a shortage of key tRNAs, we propose that the lack of fitness results from an inability to rewire the tRNA-modification profile. Altogether, our results support a modular network: loss of one node (e.g., m^7^G46) has a modest impact when other modules remain robust but becomes catastrophic when buffering modifications are weak or when codon demand focuses translation on a few vulnerable isodecoders. Resistance to METTL1 loss arises from the capacity to establish compensatory networks, largely relying on D47 to restore the tRNA elbow and coaxial stacking.

### DUS3L as a Biomarker and Mechanistic Modulator of METTL1 Sensitivity

Our model predicts that METTL1-sensitive cell lines could acquire resistance if D47 levels on tRNAs were increased. In fact, the METTL1 sensitivity across our cell-line panel inversely correlates with DUS3L abundance (hence D47), and acute METTL1 depletion by siRNA elevates DUS3L protein in OE33 and OE21 cells. However, this up-regulation is short-lived in the sensitive OE21 and absent in OE21 METTL1-hypomorphs, suggesting a cell-line–specific deleterious effect if DUS3L exceeds a certain threshold. Accordingly, DUS3L downregulation by siRNA speeds proliferation in OE21, while its overexpression proved harmful. At the opposite end, DUS3L overexpression promotes cellular fitness in OE33, and it seems to work as a “flexibility dial” at the tRNA elbow, increasing enzyme and D47 on tRNA to compensate for the lack of METTL1 activity when necessary.

The reasons for the strikingly different response to DUS3L in OE21 and OE33 await further investigation. Previous studies show that DUS3L knockout reduces protein translation and impairs cell proliferation^52^ while its overexpression predicts poor prognosis in kidney cancer, implying a context-dependent pro-growth function^53^—as observed in OE33. However, there is no indication of a general oncogenic role and, we cannot exclude that DUS3L overexpression might drive ectopic dihydrouridylation of select mRNAs, repressing their translation^54,55^ and thereby contributing to toxicity in specific cell lines, as could be the case for OE21. In any case, its sensitivity to the lack of METTL1 seems to come from the combination of its failure to invoke the DUS3L compensatory response plus a higher dependency on certain isodecoder particularly vulnerable to the lack of m^7^G46. Those include short-variable-arm (type I) tRNAs (e.g., Ala, Gly, Val, Thr, Lys) whose elbow stability leans on positions 46–47, and isodecoders with intrinsically weak acceptor/T-stem interactions (e.g. the canonical G3:U70 identity pair in all tRNA-Ala)^43,45,55,56^. Remarkably, these codons display a marked usage bias in OE21 that is further intensified by METTL1/WDR4 loss.

Recently, a catalysis-independent role for METTL1 in cell proliferation has been reported. Specifically, deletion of METTL1, but not its catalytic mutants, impaired aminoacylation by disrupting a complex with the multi-aminoacyl-tRNA synthetase (MSC). In our screen in OE33 cells, aminoacyl-tRNA synthetases did not score among significant hits. Therefore, as noted by the authors of the cited report, the m⁷G46-independent function of METTL1 in tumorigenesis may have cell type– specific relevance^57^.

Although broader studies will be needed to establish the scope of our findings, low DUS3L stratifies METTL1-sensitive cancers across our cell-line panel. Thus, DUS3L emerges as a practical, single-gene biomarker of METTL1 vulnerability that could be deployed for patient selection. In addition, our synthetic-lethal screen highlights DUS family members (DUS1L–DUS3L) as appealing targets, supporting combination strategies with METTL1 inhibitors to overcome resistance.

### Limitations of the study

This work provides, to our knowledge, the first mechanistic analysis of how cancer cells tolerate loss of METTL1 activity—a modification pathway of high therapeutic interest. We focus on the METTL1-resistant esophageal adenocarcinoma line OE33, and use the METTL1-sensitive OE21 line as a comparator.

Two caveats merit note. First, OE21 mutants are *WDR4* knockouts rather than *METTL1* directly (the latter don’t survive in the OE21 bacground). Because WDR4 is the obligate partner of METTL1, its knockout is a strong proxy for METTL1 loss; nevertheless, WDR4-specific functions independent of METTL1 cannot be completely excluded. Second, although DUS3L acts primarily on tRNAs, activity-based profiling has suggested low-stoichiometry mRNA targets, so minor contributions from non-tRNA substrates cannot be ruled out during DUS3L overexpression or depletion experiments.

Overall, the concordance between tRNA-directed modification changes, structural/functional readouts, and cellular phenotypes argues that the dominant effects we report are tRNA-mediated.

## Resource availability

### Lead contact

Requests for further information and resources should be directed to, and will be fulfilled by, the lead contact, Eric Miska

### Materials availability

All reagents are listed in the key resources table.

### Data and code availability

Raw data is available on the Short Read Archive under the project accession PRJNA1344770.

## Acknowledgments

This work was supported by CRUK (grant code: DRCRPG-Nov24/100007) and WT (grant code: 219475/Z/19/Z). We thank E. Yankova, K. Tzelepis and M. Eleftheriou for advice on the competition assay; S. Lestari for technical assistance on the SL screen; C. Livi and A. Michaelidou for support on the RNA-seq analysis; H. Fischl and O. Rausch for sharing mim-seq protocols, oligos and unpublished results; the Biophysics Facility in the Department of Biochemistry, University of Cambridge for access to instrumentation and Dr. A. Lambowitz for the generous gift of the TGIRT-III under material transfer agreement.

## FIGURE LEGENDS

**Figure S1, related to Figure 1**

A. Proliferative effect of METTL1 down regulation on the cancer cell lines shown in Figure 1A. A panel of 10 cancer cells lines were transfected with METTL1 siRNA and non-targeting scramble siRNA at 2.5nM; medium was changed after 18h and proliferation was followed by Incucyte. Down regulation of METTL1 was assessed at day 4-5 post transfection, by Western blot with anti METTL1antibody. M= MW marker. In each panel the average of the growth curves for the control (scrambled siRNA) for each cell line is normalized so that it reaches a maximum of 1 and plotted in grey. The growth curves for the METTL1 knockdowns in each cell line are then represented by averaging the coefficients of the logistic fits and computing the 95%CI of those parameters. These parameter intervals are then used to recreate the average growth curves (solid lines, coloured) and the curves that represent the 95%CI (dashed lines, coloured).

B. Boxplots of the logistic fit parameters for each of the cell lines shown as the natural logarithm of ratio of that parameter for siRNA knockdown to the scrambled control. Error bars indicate the mean and standard deviation of each distribution.

C. Characterization of METTL1 dependency by competition assay. OE21-Cas9 and OE33-Cas9 expressing GFP-tagged METTL1-targeting gRNAs (gRNA1 or gRNA2) or a non-targeting GFP control (No target) where lentiviral infected at ∼50% efficiency. The GFP⁺ population was quantified by flow cytometry every two days and normalized to day 4 post-infection (set as 100%). Data represent biological triplicates. Statistical analysis was performed using two-way ANOVA with Tukey’s multiple comparisons test (α = 0.05); *P ≤ 0.05, **P ≤ 0.01, *P ≤ 0.001. Error bars represent mean ± SD.

D. METTL1 knockdown reduces WDR4 levels in METTL1-sensitive OE21 and METTL1-resistant OE33 cells. Whole-cell lysates from ∼75% confluent cultures were transferred to nitrocellulose membrane and probed sequentially for METTL1, WDR4, and H3 (loading control) with specific antibodies.

E. Schematic of the *METTL1* cDNA showing the positions of guide RNAs, siRNAs, and the *METTL1* alleles used in this study.

F. Immunoblot analysis of OE21 rescue experiment shown in Figure 1B. Cells stable expressing empty PCMV vector (E.V) or METTL1 siRNA–resistant wild-type and catalytic mutant alleles (**t**METTL1, transfected), were transfected with siRNA targeting endogenous METTL1 (**e**METTL1, endogenous) and non-targeting scramble siRNA (scrb. siRNA). All METTL1 alleles were detected using a METTL1-specific antibody. The same membrane was re-probed with anti-Histone H3 as a loading control.

G. Chromatin immunoprecipitation (ChIP) analysis of POLR3A occupancy in OE21 and OE33 cells. Chromatin was immunoprecipitated with anti-POLR3A or anti-GFP (negative control). ChIP-qPCR shows POLR3A binding at POL III– transcribed loci (Leu-tRNA and 5S rRNA) and a non–POL III-transcribed region (negative control). Data represent three independent experiments. Statistical analysis was performed using two-way ANOVA with Tukey’s multiple comparisons test (α = 0.05); *P ≤ 0.05, **P ≤ 0.01, *P ≤ 0.001. Error bars indicate mean ± SD.

H. tRNA transcription in OE33 *versus* OE21. POLR3A ChIP–seq occupancy in OE21 and OE33 cells are displayed at (I) loci, (II) isodecoder, (III) isoacceptor, and

(IV) isotype levels. The results represent 3 independent experiments (n=3).

I. m7G46 methylome in OE33 *versus* OE21. Misincorporation rates at G46 detected by tRNA-mim-seq in OE21 and OE33 cells are displayed at (I) isodecoder,

G. (II) isoacceptor, and (III) isotype levels. The results represent 3 temporal independent experiments (n=3).

**Figure S2, related to Figure 2**

A. Generation of *METTL1* knockout OE33 single-cell clones. Schematic of the cloning workflow. A wild-type clone WT/WT generated by “non-targeting gRNA” (OE33_WT_) and three homozygous *METTL1* knockout clones (METTL1 KO1–KO3) were selected for downstream studies. Genotypes were confirmed by TIDE analysis.

B. Proliferation rates of individual OE33 *METTL1* knockout clones. Cells were seeded at ∼10% confluency, and growth was monitored every 3 hours using Incucyte. Plots show the proliferative rate (log Δr) normalized to the OE33 wild-type single-cell clone. Each dot represents a biological replicate. Error bars represent the 95% confidence interval of the mean, when bars are overlapping, the distributions are not significantly different at an alpha=0.05

C. Generation of *WDR4* knockout OE21 single-cell clones. Schematic of the cloning workflow. A wild-type clone WT/WT generated by infecting a “non-targeting gRNA” (OE21_WT_) and four homozygous *WDR4* knockout clones (*WDR4* KO1– KO4) were selected for downstream studies. Genotypes were confirmed by TIDE analysis.

D. Proliferation rates of individual OE21 *WDR4* knockout clones. Cells were seeded at ∼10% confluency, and growth was monitored every 3 hours using Incucyte. Plots show the proliferative rate (log Δr) normalized to the OE21 wild-type single-cell clone. Each dot represents a biological replicate. Error bars represent the 95% confidence interval of the mean, when bars are overlapping, the distributions are not significantly different at an alpha=0.05.

E. OE33 Northern blot analysis of tRNA. Total RNA from parental, WT and METTL1 KO single cell clones (3 micrograms) were resolved in TBE-Urea gels stained with Sybergold (left panels), transferred to Nylon membranes and hybridized, first with oligos against tRNAs (as indicated in middle panels) and secondly with an oligo against the POLIII transcript ribosomal 5S, used as additional loading control. tRNA are marked by a red asterisk (**\***), r5S is marked by a purple asterisk (**\***).

F. 21 Northern blot analysis of tRNA. Total RNA from parental, WT and *WDR4* KO single cell clones (3 micrograms) were resolved in TBE-Urea gels stained with Sybergold (left panels), transferred to Nylon membranes and hybridized, first with oligos against tRNAs (as indicated in middle panels) and secondly with an oligo against the POLIII transcript ribosomal 5S, used as additional loading control. tRNA are marked by a red asterisk (**\***), r5S is marked by a purple asterisk (**\***).

**Figure S4, related to Figure 4**

A. Schematic illustrating the synthetic lethal screen in wild-type OE33_WT_ and *METTL1* KOs clones.

B. Genome-wide Synthetic Lethality Screen Ranked by Gene Effect Scores. Genes are ordered along the x-axis from strongest synthetic lethal interactions (left) to weakest (right). Each point represents a gene, plotted according to its rank. Dropouts (negative beta scores) are highlighted in blue, while positively selected genes (positive beta scores) are highlighted in pink. Left: *METTL1* KO1 *versus* OE33_WT_ clone; right: *METTL1* KO2 *versus* OE33_WT_ clone.

C. Dotplot showing significantly enriched terms identified from MAGeCKFlute analysis using the hypergeometric test (HGT) of negatively selected genes in *METTL1* KO1 (top) and *METTL1* KO2 (bottom). The x-axis represents the – log10(FDR) of enrichment, and the y-axis shows the top enriched pathways or terms ranked by statistical significance. The size of the dots indicates the Net Enrichment Score (NES).

D. Immunoblot analysis of the proliferation assays shown in Figure 3D. Cells were collected 5 days after transfection with siRNA targeting each of the hits from the SL screen (panels from top to bottom: XPOT-siRNA, NSUN2-siRNA, DUS1L-siRNA, DUS2L-siRNA and DUS3L-siRNA) or a non-targeting scramble siRNA as control (scrb. siRNA). 45 micrograms of protein extracts from each clone were separated in SDS-PAGE gels, transferred to Nitrocellulose membranes and blotted firstly with specific antibody against each SL target (as indicated) and re-probed with anti-GAPDH as a loading control.

**Figure S5, related to Figure 5**

A. Thermal Denaturation Profiles of Modified and Unmodified tRNA Monitored by Circular Dichroism at 270 nm. Thermal melting curves of Tyr-GTA-tRNA carrying different combinations of post-transcriptional modifications were recorded by monitoring ellipticity at 270 nm as a function of temperature. Data are shown for tRNA oligos containing D16D17D20m^7^G46D47 (blue) or unmodified tRNA (grey). Samples were normalized to initial ellipticity values to allow direct comparison. Error bars represent the standard deviation from 2 independent replicates.

**Figure S6, related to Figure 6**

A. tRNA modification map for parental OE33 and OE21 cells and their derivatives. Heatmap representing changes in m^2,2^G26, m^7^G46, D47, and m^1^A58, inferred from reverse-transcription misincorporation and deep sequencing (tRNA mim-seq). tRNA isotypes are grouped on the left; isodecoders are listed on the right. In blue WT: OE33_P_ plus OE33_WT_, KO: *METTL1*KO1 plus *METTL1*KO2; In green WT: OE21_P_ plus OE21_WT_, KO: *WDR4*KO2 plus *WDR4*KO3.

B. Quantification of DUS3L mRNA in parental OE33_P_, wild-type OE33_WT_ and *METTL1*KO clones. Total RNA from the samples in B was used to synthesize cDNA and the levels of DUS3L were assessed by quantitative PCR with specific primers. The plot shows the data of 3 independent experiments, internally normalized to GAPDH.

C. Immunoblot analysis of DUS3L protein levels from samples described in B. Total protein extracts (45 micrograms) were transferred to Nitrocellulose membrane and immunoblotted firstly with anti-DUS3L and sequentially with anti-GAPDH as a loading control.

D. Immunoblot analysis of DUS3L protein levels in parental, OE21_P_, wild-type OE21_WT_ and *WDR4* KO1-3 clones. Total protein extracts (45 micrograms) were transferred to Nitrocellulose membrane and immunoblotted firstly with anti-DUS3L and sequentially with anti-H3 as a loading control.

**Figure S7, related to Figure 7**

A. DUS-L family mRNA expression in parental OE33_P_ (resistant) and OE21_P_ (sensitive) cell lines. Poly(A)+ RNA-seq libraries were prepared from 3 µg total RNA (n = 3 biological replicates per line). Box plots summarize library-normalized expression of dihydrouridine synthase–like (DUS-L) genes.

B. METTL1 CRISPR dependency *versus* DUS3L expression across the panel of Cancer Cell Lines shown in Figure 1A.

**Figure 7. METTL1 resistance correlates with high DUS3L protein expression.**

**I.**Immunoblot analysis of DUS3L levels in OE33_P_ and OE21_P_ parental cell lines. Total protein extracts (45 micrograms) from cells grown at 50% and 100% confluency, were transferred to Nitrocellulose membrane and immunoblotted firstly with anti-DUS3L and sequentially with anti-METTL1 and anti-GAPDH as a loading control.

**J.**Immunoblot analysis of DUS3L levels across the panel of Cancer Cell Lines shown in Figure 1A. Total protein extracts (45 micrograms) from cells grown at 75% confluency, were transferred to Nitrocellulose membrane and immunoblotted firstly with anti-DUS3L and sequentially with anti-METTL1 and anti-GAPDH as a loading control.

**K.**Immunoblot analysis of endogenous DUS3L and overexpressed myc-DUS3L in parental OE21. Total protein extracts (45 micrograms) from stable multiclonal population generated by transfection of empty-pCMV vector or pCMV-myc-DUS3L plasmids, were transferred to Nitrocellulose membrane and immunoblotted firstly with anti-DUS3L (note signal acquired after 37 sec exposure) and sequentially anti-H3 as a loading control. Endogenous DUS3L is marked with red asterisk (**\***), transfected myc-DUS3L is marked with purple asterisk (**\***)

**L.**Representative light-microscopy images of OE21 cells with endogenous DUS3L (empty-vector control, pCMV; top) or DUS3L overexpression (pCMV-myc-DUS3L; bottom). Images were acquired 4 days after completion of antibiotic selection of transfected cells

**M.**Immunoblot analysis of endogenous and pCMV-overexpressed DUS3L in parental OE33. Total protein extracts (45 micrograms) from stable clones generated by transfection of empty-pCMV vector or pCMV-myc-DUS3L overexpression plasmid cells, were transferred to Nitrocellulose membrane and immunoblotted firstly with anti-DUS3L (note signal acquired after 5 sec exposure) and sequentially anti-H3 as a loading control. Endogenous DUS3L is marked with red asterisk (**\***), transfected myc-DUS3L is marked with purple asterisk (**\***)

**N.**Representative light-microscopy images of OE33 cells with endogenous DUS3L (empty-vector control, pCMV; top) or DUS3L overexpression (pCMV-myc-DUS3L; bottom). Images were acquired 4 days after completion of antibiotic selection of transfected cells.

**O.**Proliferation rates of parental OE33 stable clones expressing either endogenous (empty-pCMV, grey) overexpressed DUS3L levels (pCMV-myc-DUS3L, blue), as in panel E. Cells were seeded at ∼10% confluency, and growth was monitored every 3 hours using Incucyte. (n=6)

**P.**Proliferation rates of parental OE21 cells expressing transfected with either scramble control siRNA (grey) or DUS3L siRNA (green). Cells were seeded at ∼10% confluency, and growth was monitored every 3 hours using Incucyte. (n=3)

## Notes

### Competing Interest Statement

The authors have declared no competing interest.

