## Supplementary figures and images for "Compensatory tRNA Modification by DUS3L Confers Resistance to METTL1 Loss in Oesophageal Cancer"

### S1

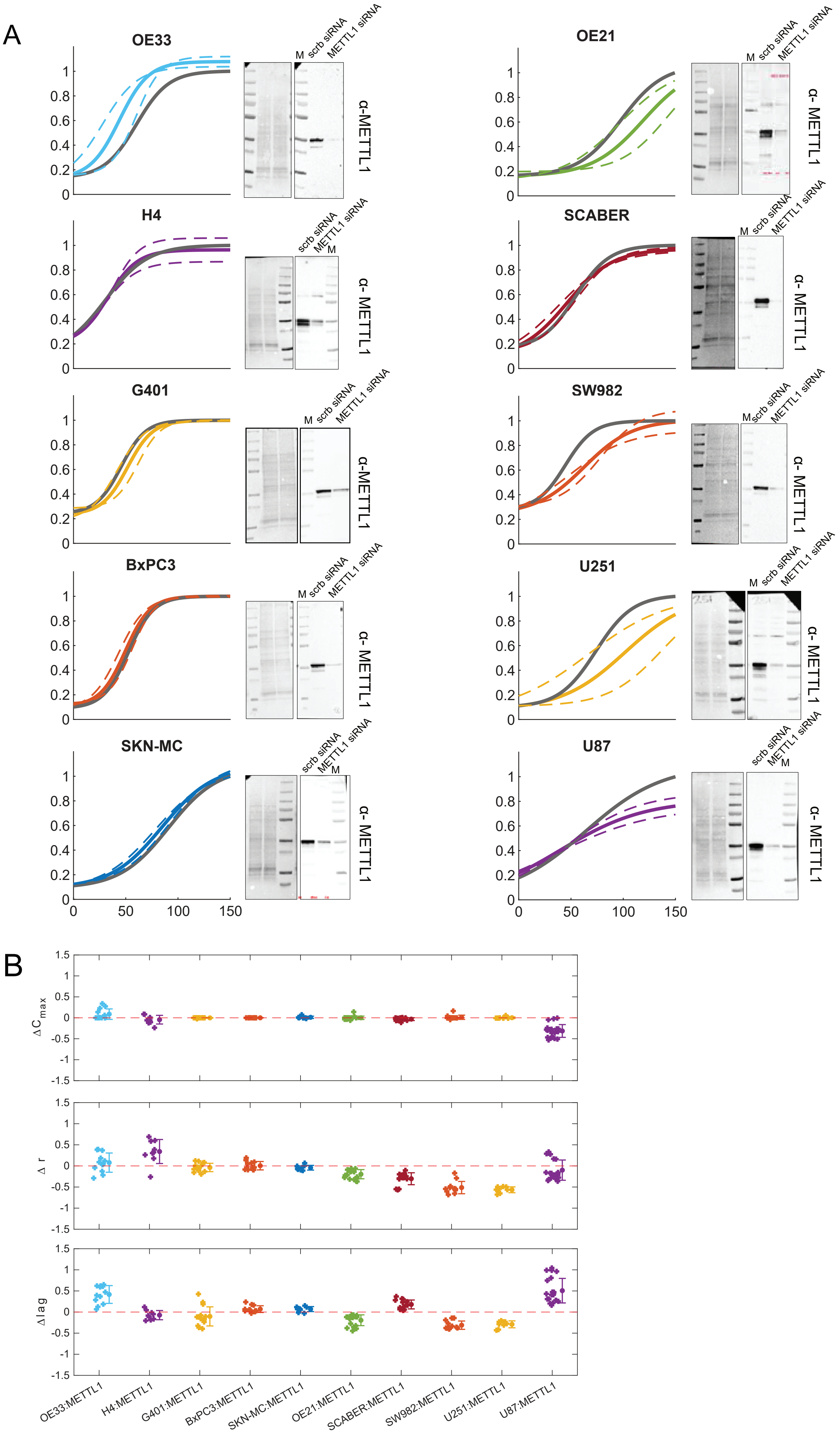

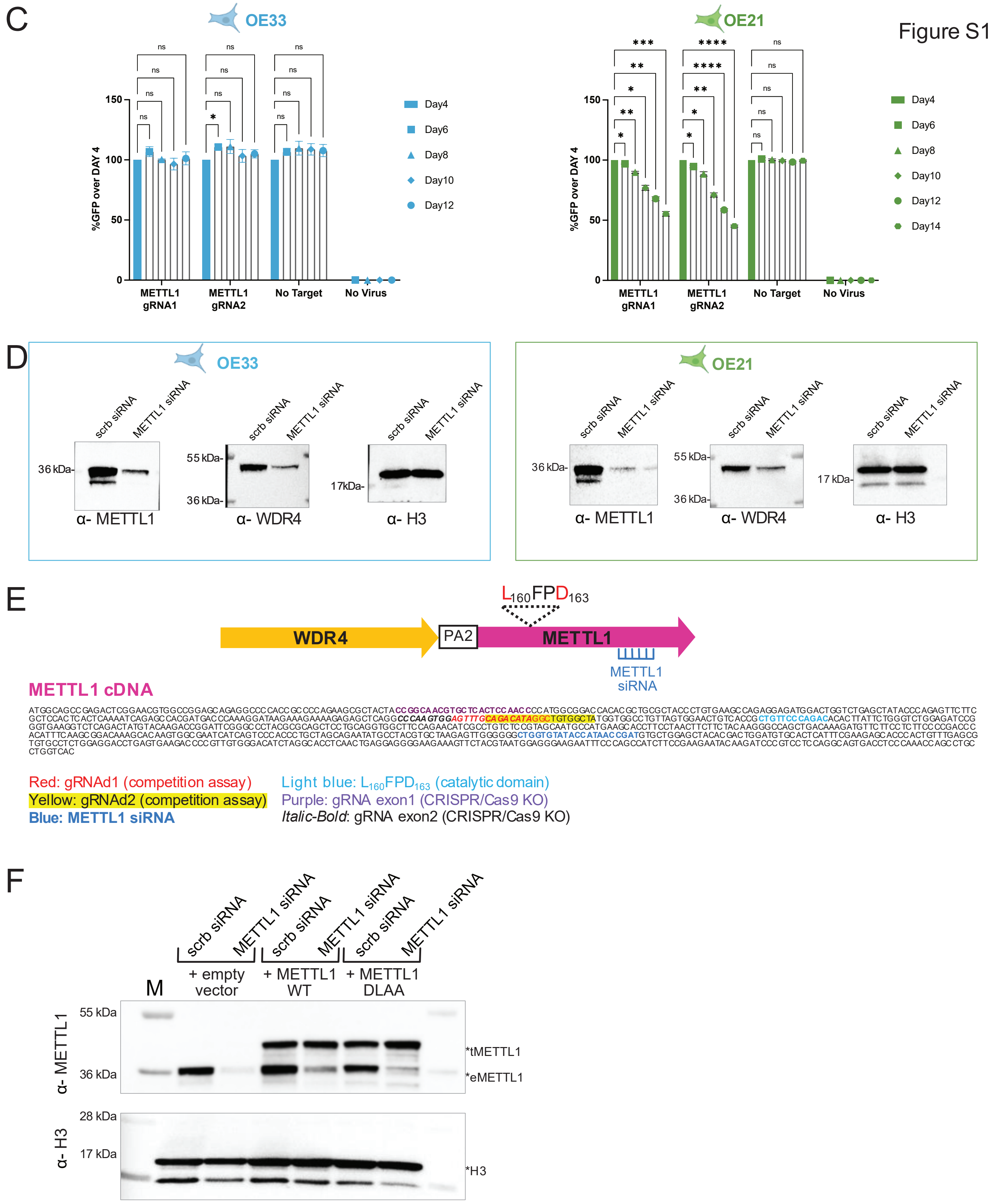

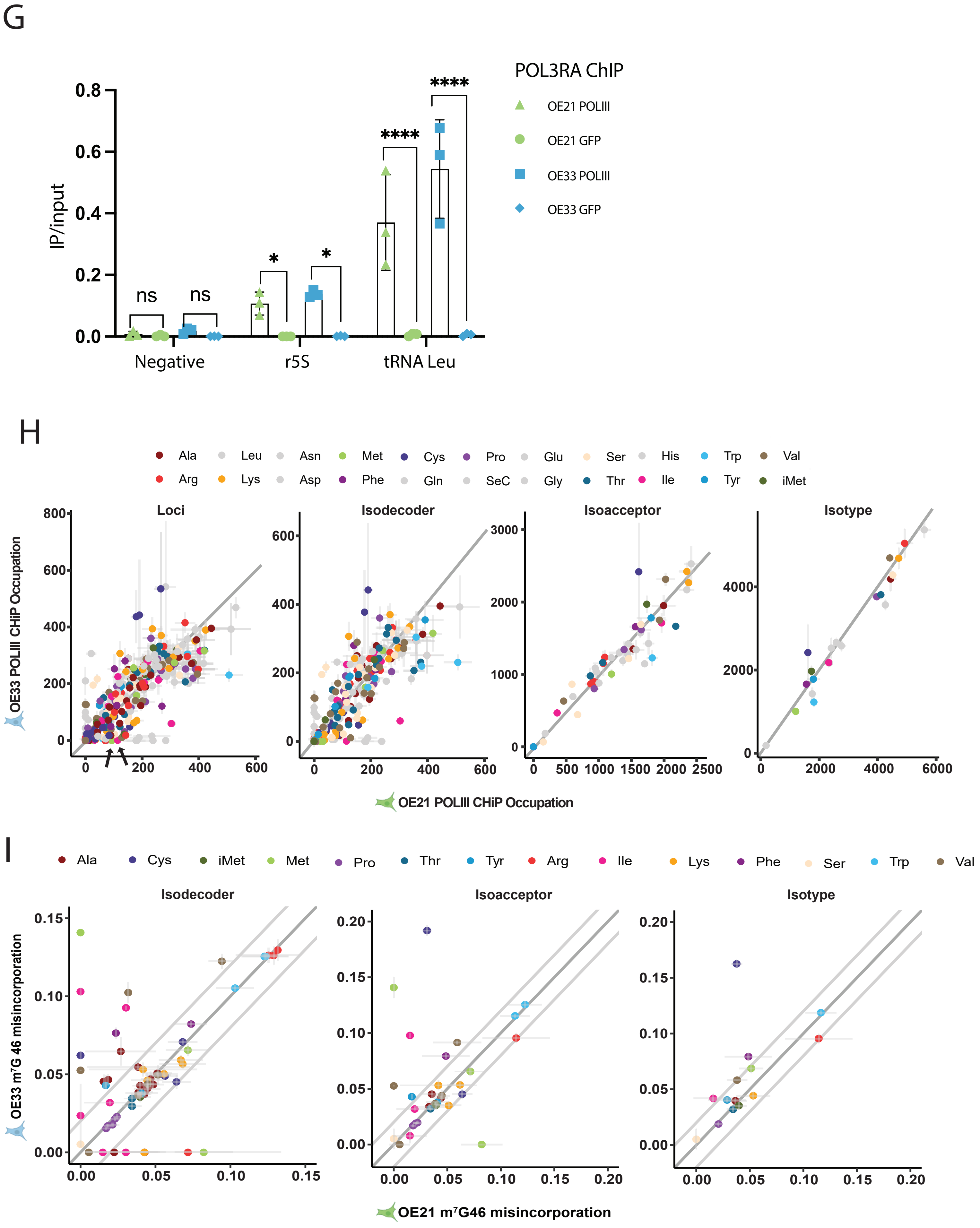

### S2

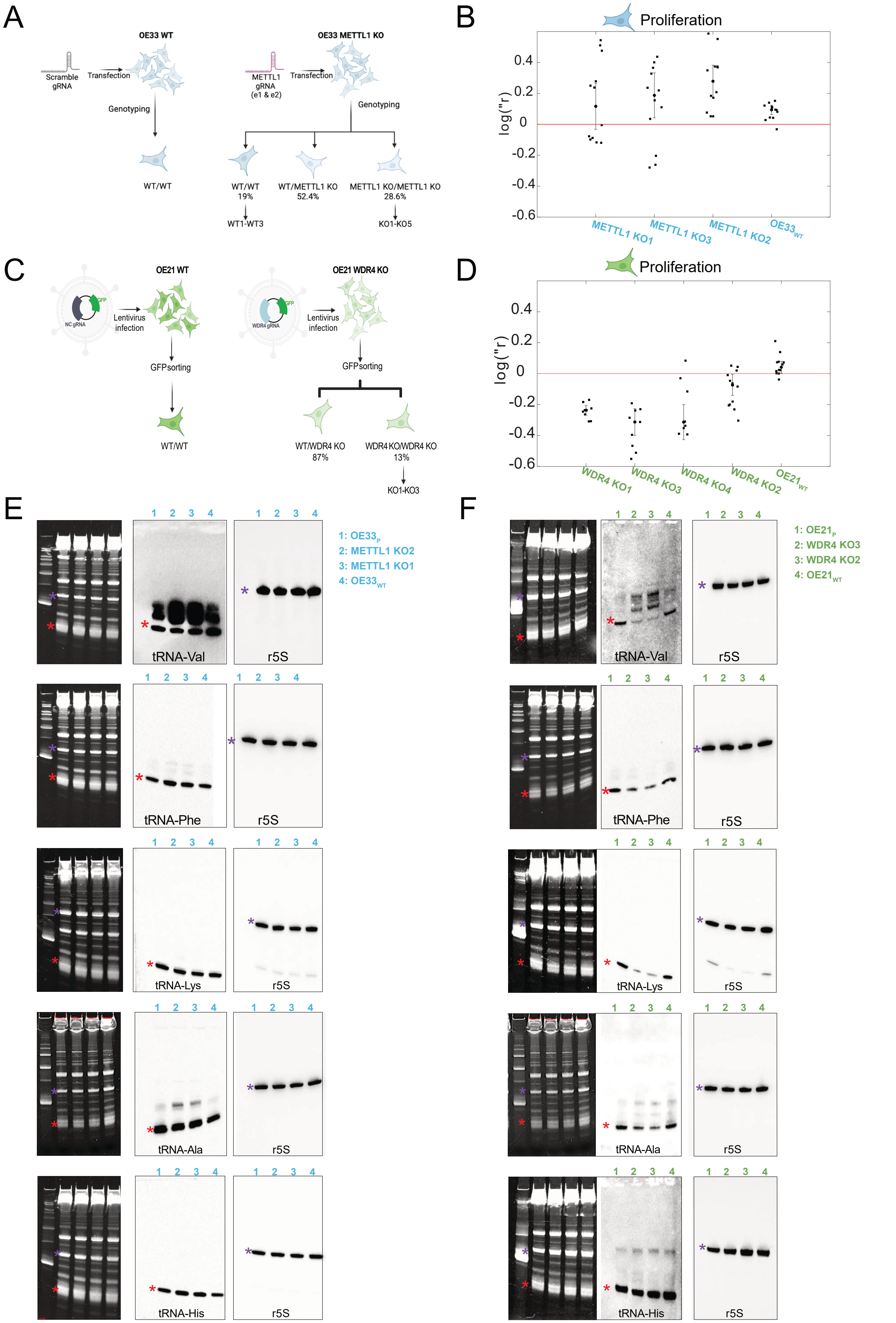

### S4

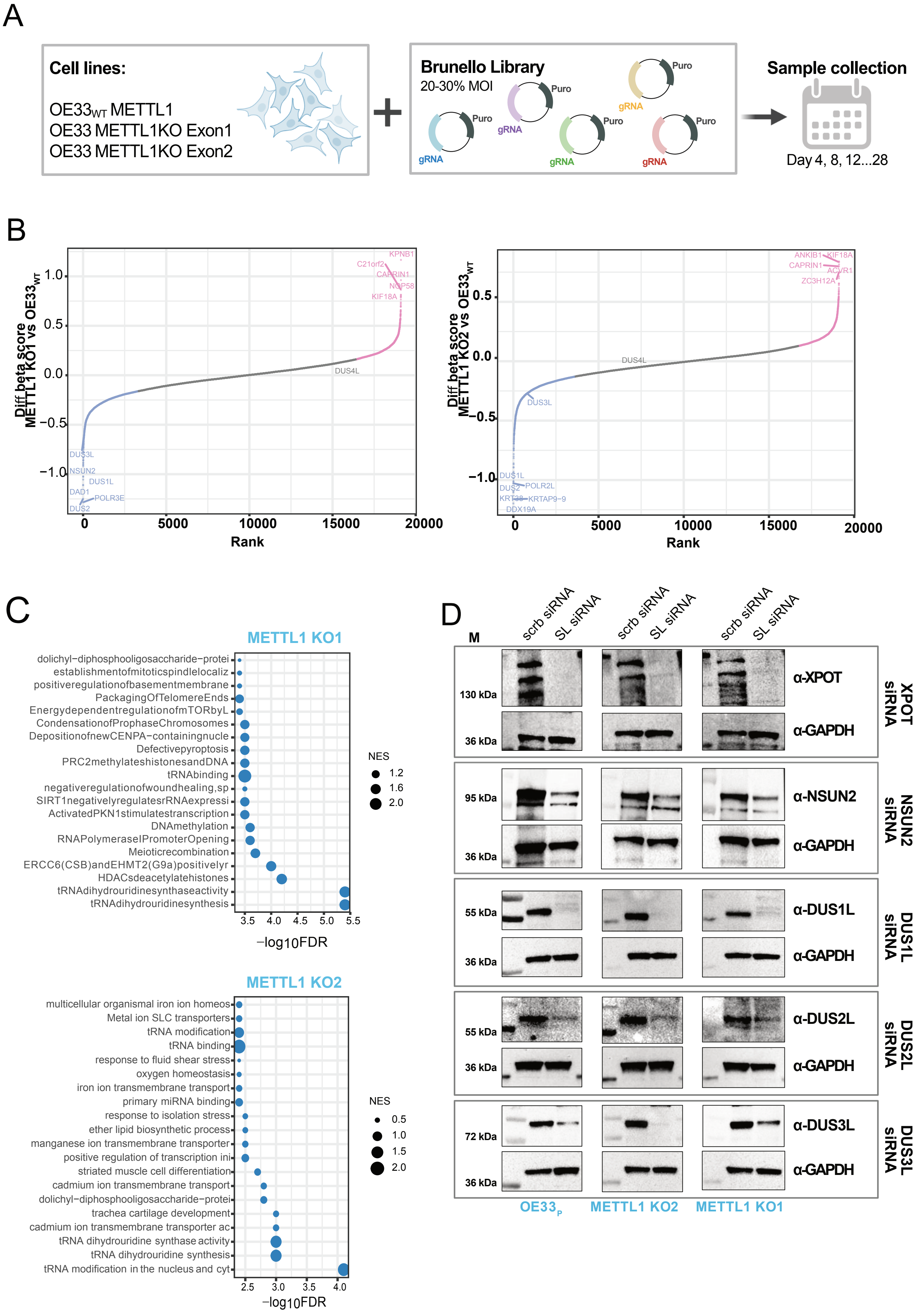

### S6

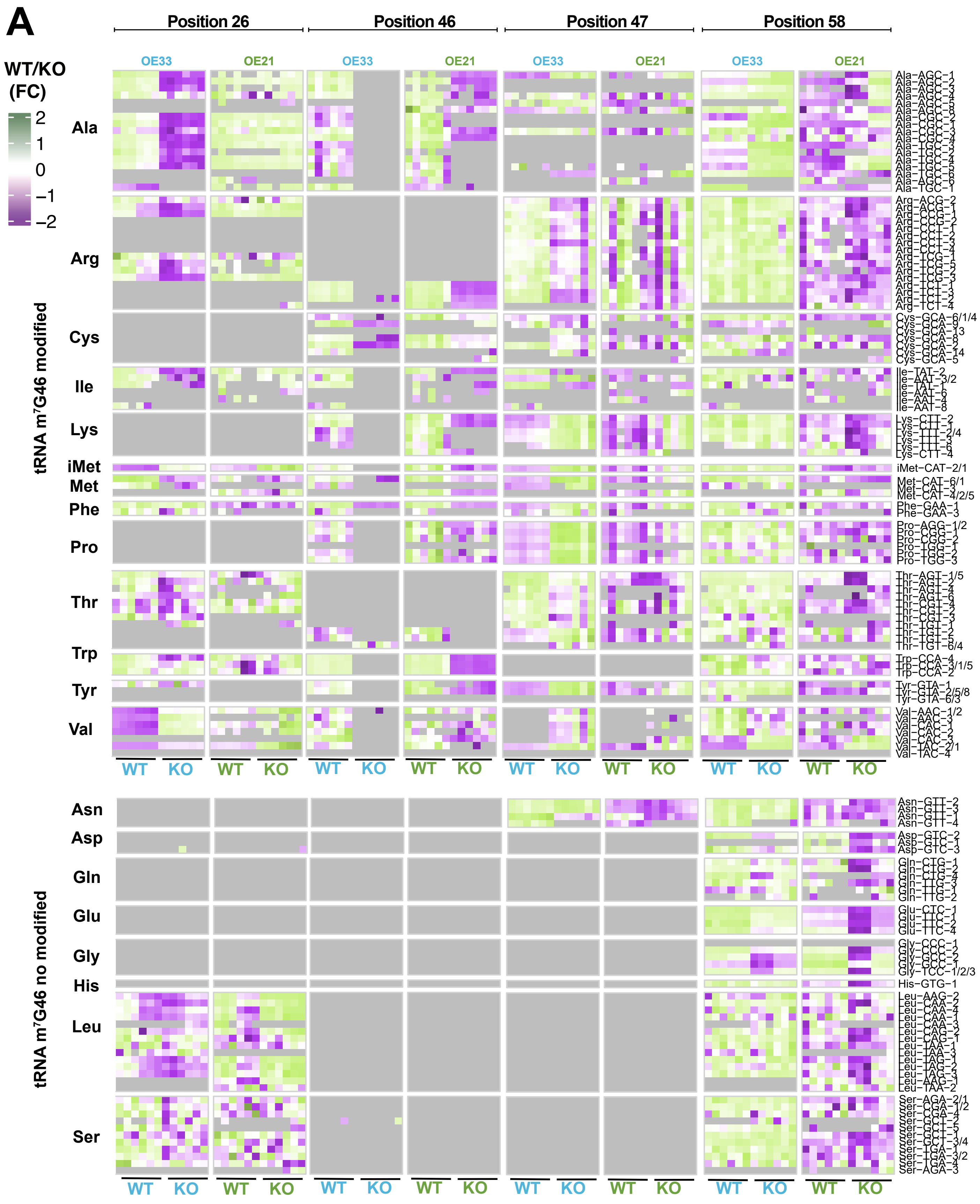

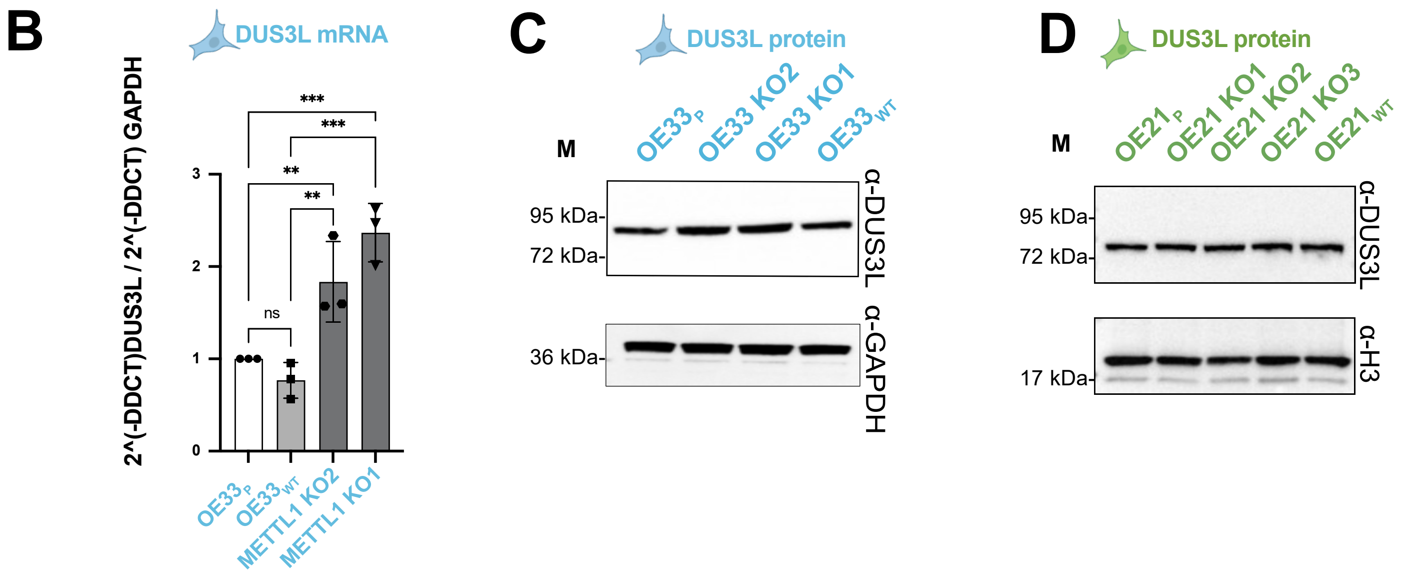

### S7

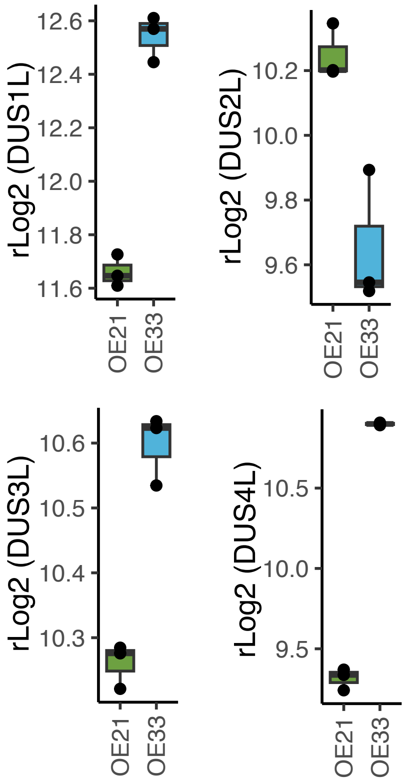

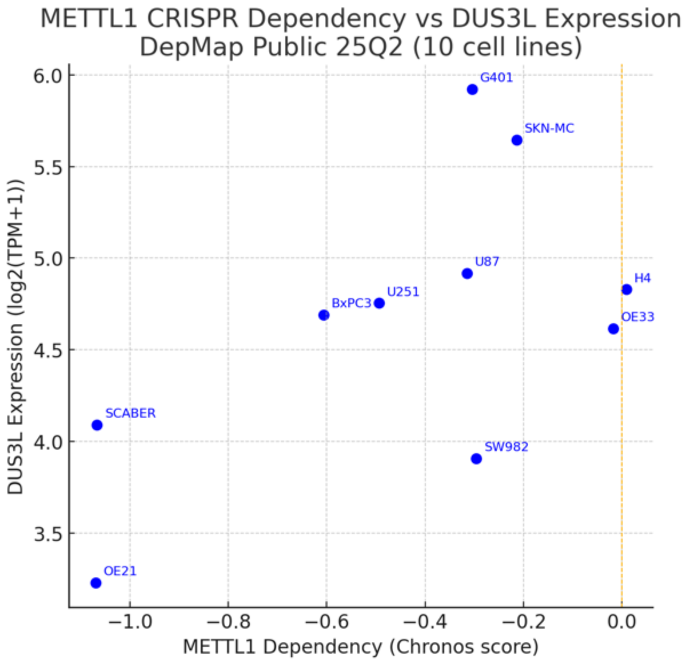
