## Supplementary material for "Compensatory tRNA Modification by DUS3L Confers Resistance to METTL1 Loss in Oesophageal Cancer": S5

A

>Unmodified
CCUUCGAUAGCUCAGUUGGUAGAGCGGAGGACUGUAGAUCC
UUAGGUCGCUGGUUCGAAUCCGGCUCGAAGGA

>D16 D17 D20 m G46 D47
CCUUCGAUAGCUCAGDDGGDAGAGCGGAGGACUGUAGAUCC
UUAGGDCGCUGGUUCGAAUCCGGCUCGAAGGA

DUS1L-deposited Dihydrouridine
DUS2L-deposited Dihydrouridine
DUS3L-deposited Dihydrouridine
METTL1-deposited N7-methylguanosine

B

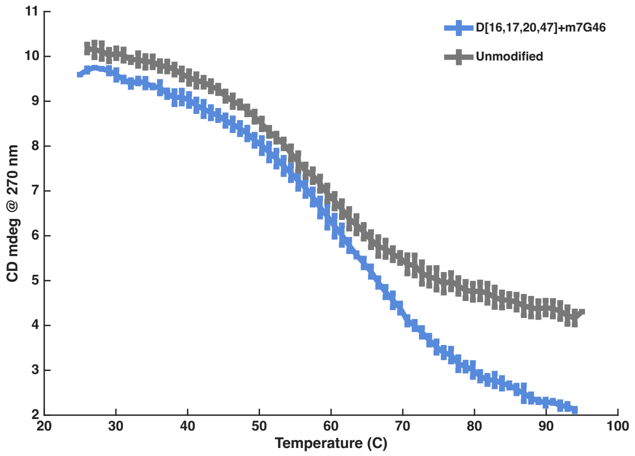
